# Spatial mapping of immunosuppressive CAF gene signatures in H&E-stained images using additive multiple instance learning

**DOI:** 10.1101/2024.08.12.607604

**Authors:** Miles Markey, Juhyun Kim, Zvi Goldstein, Ylaine Gerardin, Jacqueline Brosnan-Cashman, Syed Ashar Javed, Dinkar Juyal, Harshith Pagidela, Limin Yu, Bahar Rahsepar, John Abel, Stephanie Hennek, Archit Khosla, Amaro Taylor-Weiner, Chintan Parmar

## Abstract

The relative abundance of cancer-associated fibroblast (CAF) subtypes influences a tumor’s response to treatment, especially immunotherapy. However, the extent to which the underlying tumor composition associates with CAF subtype-specific gene expression is unclear. Here, we describe an interpretable machine learning (ML) approach, additive multiple instance learning (aMIL), to predict bulk gene expression signatures from H&E-stained whole slide images (WSI), focusing on an immunosuppressive LRRC15+ CAF-enriched TGFβ-CAF signature. aMIL models accurately predicted TGFβ-CAF across various cancer types. Tissue regions contributing most highly to slide-level predictions of TGFβ-CAF were evaluated by ML models characterizing spatial distributions of diverse cell and tissue types, stromal subtypes, and nuclear morphology. In breast cancer, regions contributing most to TGFβ-CAF-high predictions (“excitatory”) were localized to cancer stroma with high fibroblast density and mature collagen fibers. Regions contributing most to TGFβ-CAF-low predictions (“inhibitory”) were localized to cancer epithelium and densely inflamed stroma. Fibroblast and lymphocyte nuclear morphology also differed between excitatory and inhibitory regions. Thus, aMIL enables a data-driven link between tissue phenotype and transcription, offering biological interpretability beyond typical black-box models.

## Introduction

As a tumor develops, a complex interplay between the cancer cells and the surrounding stroma, or tumor microenvironment (TME), strongly influences disease progression. The TME contains a number of non-cancerous cell types, including immune cells, endothelial cells, and cancer-associated fibroblasts (CAFs).

CAFs are cells of mesenchymal origin that reside within a tumor and are non-epithelial, non-endothelial, non-immune, and non-cancerous. CAFs have been widely studied, revealing a range of both tumor-suppressing and tumor-promoting functions involving remodeling of the extracellular matrix (ECM), metabolite secretion, and regulation of anti-tumor immunity ^1^. In recent years, the heterogeneity of CAFs has been increasingly appreciated, and CAF subpopulations have been identified with distinct gene expression profiles and functions^2,3^. Importantly, several subsets of CAFs are known to function in an immunosuppressive manner through the upregulation of inhibitory immune modulators (e.g., PD-L1) directly on the CAFs or through stimulating upregulation of such molecules in cancer cells through the secretion of cytokines such as CXCL5 and TGFβ. Examples of such CAFs include the myCAF^4^ and CAF-S1^5,6^ subpopulations.

In particular, recent work has identified that CAFs expressing LRRC15 are found in many cancer subtypes and are associated with immunosuppressive activity and poor response to immunotherapy in pancreatic, bladder, breast, lung, renal, and head and neck cancer^7–10^. LRRC15, itself, is activated by transforming growth factor-beta (TGFβ)^7^, suggesting a functional role for this signaling network in the maintenance of this CAF population. Interestingly, the LRRC15 gene is also a component of the ecm-myCAF gene signature in breast cancer^10^. This CAF subpopulation, characterized by a gene signature relating to collagen synthesis, has been shown to be associated with an immunosuppressive microenvironment marked by few CD8+ lymphocytes and enrichment of regulatory T cells and PD-1/CTLA-4 CD4+ lymphocytes. Furthermore, regulatory T cells were found to convert ecm-myCAFs into a related CAF population (TGFβ-myCAF). TGFβ-myCAFs, characterized by a gene signature indicative of TGFβ stimulation, were associated with ecm-myCAFs and the same immunosuppressive features. Notably, in patients with melanoma and NSCLC treated with checkpoint inhibitors, gene expression signatures (GES) indicative of ecm-myCAF and TGFβ-myCAF populations were enriched in non-responders^10^.

Despite the clear relationship between CAFs and outcome in many cancers, CAFs have proven to be unreliable as a histopathological biomarker, given that there is no specific marker for this heterogeneous cell type^3^. As such, Dominguez and colleagues identified a gene signature (“TGFβ-CAF”) associated with the presence of LRRC15+ CAFs^8^. However, despite increased research into the applicability of gene signatures to cancer in the clinical setting, these proposed biomarkers have not been widely adopted in the clinic. While the cost for performing sequencing analyses has, generally, decreased, these assays remain time consuming (especially relating to data analysis), costly, and require considerable high-quality tissue, hindering the widespread clinical adoption of transcriptomic biomarkers^11,12^.

Conversely, hematoxylin and eosin (H&E) staining of tumor biopsies is a routine and nearly universal process. The scanning of H&E slides can generate digital whole slide images (WSI) with a wealth of information, on which machine learning (ML) algorithms can be deployed to predict diagnosis, prognosis, tumor grade, treatment response, and recurrence^13,14^. In addition, these models can be utilized to infer additional relevant information about the tissue sample, including cell and tissue composition, biomarker expression, and gene expression patterns. That said, WSIs are extremely information dense, with each image containing billions of pixels and thousands of cells and tissue regions. The potential for extracting and inferring information from these images is great, but challenges exist due to WSI size. Furthermore, while other studies have shown the ability to predict gene expression scores from WSIs, these studies lack biological interpretability. To overcome these challenges, we developed a framework termed additive multiple instance learning (aMIL)^15^, which does not necessitate the collection of patch-level labels, allows end-to-end training on a WSI, provides the exact contribution of each image patch to a slide-level label, and allows each patch-level contribution to be visualized as a heatmap.

Here, we constructed an aMIL model to predict the TGFβ-CAF gene signature in six cancer types (BRCA, LUAD, LUSC, PRAD, BLCA, and STAD). In addition, we show that this model output is highly interpretable, allowing 1) assessment of the contribution of each patch to slide-level TGFβ-CAF predictions, 2) interrogation of the cell and tissue features associated with TGFβ-CAF expression via subsequent application of multiple additional models. The ability to conduct detailed biological inspection of our model provides the opportunity not only for validation of its function, but also to gain new insights about relevant gene expression profiles in a spatially interpretable manner.

## Methods

### Data

Cases from The Cancer Genome Atlas (TCGA)^16^ were selected from the following tumor types: bladder cancer (BLCA), breast cancer (BRCA), gastric cancer (STAD), lung adenocarcinoma (LUAD), lung squamous cell carcinoma (LUSC), and prostate adenocarcinoma (PRAD). Dataset characteristics are listed in Supplementary Table 1. Cases were selected based on availability of WSI derived from H&E-stained slides derived from formalin-fixed, paraffin-embedded sections and associated transcriptomic data. Slides were excluded if they either possessed less than 1 mm^2^ of usable tissue as defined by ArtifactDetect (PathAI, Boston, MA), or if they were flagged as “not-evaluable” based on manual pathologist QC. Each indication was split into training (60%), validation (20%) and testing (20%) datasets based on demographic, clinical, and sampling variables and CaseIDs such that all slides from a given case were grouped into the same sub-set (train/val/test). Slide level TGFβ-CAF scores were determined according to the method described by Krishnamurty and colleagues^9^ based on batch-corrected normalized RNA-Seq data acquired from the Genomic Data Commons (GDC)-processed TCGA cohorts (version 2016-12-29) from the UCSC Xena data portal^17^. Classification of slides as TGFβ-CAF Low or TGFβ-CAF High was performed using the approach described by Dominguez and colleagues, whereby the median TGFβ-CAF score from each indication’s training dataset was used to dichotomize the labels as either “Low” (< median) or “High” (≥ median)^8^. The median values for each indication, used as the cutoffs for binarization, are specified in Supplementary Table 2.

### TME Model Architecture and Training

For each cancer type examined, indication-specific PathExplore^TM^ models (PathAI, Boston, MA) were deployed to predict tissue regions and cell types in H&E-stained WSI^18^; PathExplore is for research use only and is not for use in diagnostic procedures.

Cancer epithelial cells, fibroblasts, macrophages, lymphocytes and plasma cells were predicted for all cancer types, while additional cell classes were predicted in Gastric (neutrophils and eosinophils), LUAD (granulocytes and normal cells) and PRAD (smooth muscle cells, endothelial cells, and normal epithelial cells). Model performance for the prediction of cell types was assessed by comparing model predictions to pathologist annotations in nested pairwise fashion ^19^. Model performance metrics for BRCA, LUAD, LUSC, and PRAD cell predictions were previously reported by Abel and colleagues^18^, while performance metrics for BLCA and STAD cell predictions are shown in Supplementary Figures S1-S2 and Supplementary Tables S3 and S4. To enable comparative analysis across all indications, five main cell classes (cancer epithelial cells, fibroblasts, macrophages, lymphocytes, and plasma cells) were used for analyses herein.

For all cancer types, PathExplore tissue models predicted regions of normal tissue, cancer epithelium, cancer stroma, and necrosis. Additional tissue regions were predicted for BRCA (DCIS, lymph nodes), and Gastric (Mucin). Model performance for the prediction of tissue regions was assessed by comparing model predictions to pathologist annotations either qualitatively or in nested pairwise fashion^19^. Performance metrics for BLCA, BRCA, STAD, LUAD/LUSC, and PRAD are shown in Supplementary Figure S3-S7 and Supplementary Table 5, respectively. Regions predicted to be cancer epithelium and cancer stroma were used for analyses herein.

### aMIL Model Architecture and Training

For each indication, training and validation WSI were sampled into 256×256-pixel patches at 0.5 microns per pixel resolution from regions of cancer and cancer stroma, classified by PathExplore models, for model training. Additive multiple-instance learning (aMIL) models^15^ were trained to predict the slide-level binarized TGFβ-CAF status (“high” or “low”) using these indication-specific sample sets. These models were trained for 10000 iterations with a bag size of 24 and training batch size of 32 and evaluated every 50 iterations on the validation set using the same batch and bag sizes. After training, the best performing model iteration was selected based on validation F1 score and evaluated on the test set. Receiver operating characteristic curves (ROCs) were used to evaluate the performance of the binary classifier. For other cut-point based performance metrics (e.g., accuracy, sensitivity, specificity) the optimal cut-point was defined using Youden’s J statistic ^20^ in the validation cohort and applied to evaluate the model performance in the test set.

### Stromal subtyping model architecture and training

A convolutional neural network-based model was developed to classify cancer associated stroma in H&E-stained WSI as immature, mature, densely inflamed, densely fibroblastic, or elastosis^21^. This model was trained using manual pathologist-derived annotations (N=3019) of H&E-stained whole slide images (WSIs) of PDAC obtained from the TCGA (N=126). Model performance was assessed by qualitative review by expert pathologists.

### Model Deployment and Feature Extraction

The indication-specific aMIL TGFβ-CAF models were deployed on validation and test sets of H&E stained WSI from each indication as shown in Figure 1A. At inference, aMIL heatmaps were used to generate spatial credits to derive a unique patch-level TGFβ-CAF contribution score. From the patch-level contribution score distribution of the validation set, all the patches within the top 25% of scores and bottom 25% of scores were defined as “high contribution” patches, termed “excitatory” and “inhibitory,” respectively. These high contribution patches were then selected in an aMIL-heatmap by filtering out remaining patches, leaving only excitatory or inhibitory regions. At the end of this step, slides had only high contribution patches remaining in the aMIL heatmaps (Figure 1B). For feature comparison, the aMIL heatmap was further masked to include only excitatory regions for slides predicted to be TGFβ-CAF-high, and only inhibitory regions in slides predicted to be TGFβ-CAF-low. These filtered aMIL heatmaps were then used to mask each slide’s corresponding cell, tissue, stromal subtype and nuclei heatmaps. Human interpretable features (HIFs) were extracted from aggregated excitatory or inhibitory regions on each slide. HIFs characterized the spatial distribution of cell, tissue, stromal subtype and nuclei as well as nuclear morphology within these “high contribution” regions. HIFs describing the TME (N=297) were extracted describing the relative area of tissue regions, relative count of cells, cell densities, and cell density ratios. HIFs describing nuclear morphology (N=480) were extracted describing the size, shape, texture, and stain intensity of nuclei in a cell-specific manner (e.g., nucleus area in lymphocytes). Of these features, a subset (N=54 TME HIFs and N=230 nuclear HIFs) were chosen due to their simpler biological interpretation and were included in the final analysis. Thus, the extracted HIFs quantify the biology of the WSIs pertinent to making the TGFβ-CAF classification prediction.

**Figure 1.**
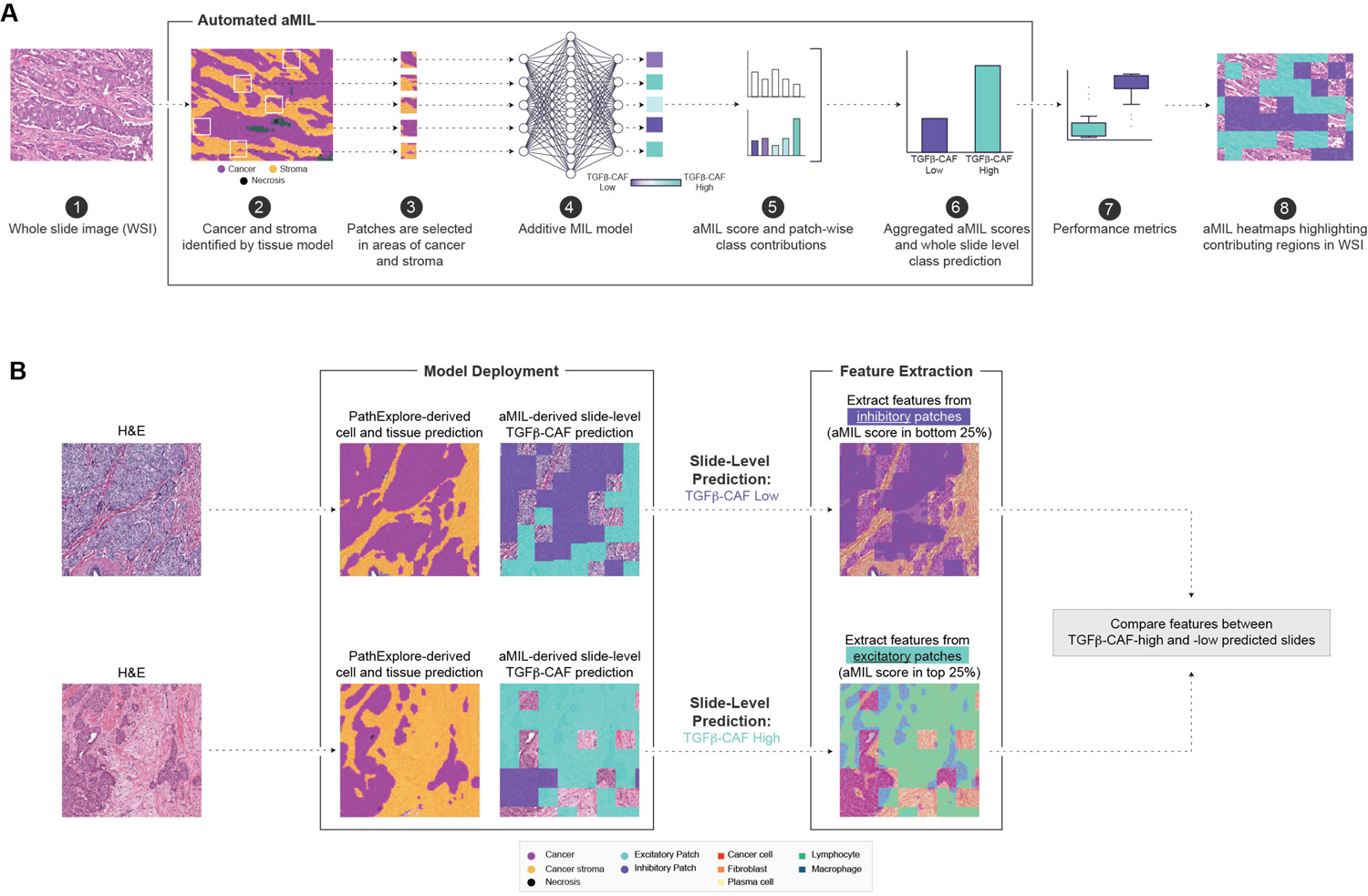
Study design. A) Schematic representation of aMIL model to predict TGFβ-CAF levels from H&E-stained WSI. Patches sampled from cancer epithelial and stroma regions of H&E WSI were fed as input to train the aMIL model to predict the slide-level binarized TGFβ-CAF status. At inference, aMIL heatmaps were used to generate spatial credits to derive a unique patch-level TGFβ-CAF contribution score (step 8). B) Approach to interpret aMIL model predictions based on spatial composition of the TME: Cell types and tissue substances were identified and superimposed with patch-wise aMIL attention scores. Features from high contribution regions (top 25th percentile of aMIL scores for predicted TGFβ-CAF high slides; bottom 25th percentile for TGFβ-CAF low slides) were extracted for further analysis.

### Statistical Analysis

We performed the Mann-Whitney U test to assess the distribution differences between TGFβ-CAF-high and TGFβ-CAF-low predicted slides for each feature. P-values were corrected for multiple testing using the Benjamini-Hochberg method. Principal component analysis (PCA) was performed on the BRCA dataset after z-score transformation. For pan-cancer analysis, clustering was performed using the agglomerative hierarchical clustering method with average linkage and Euclidean distance. Statsmodels and scikit-learn python packages were used for the statistical analyses.

## Results

### Performance of aMIL model in predicting TGFβ-CAF levels in multiple cancer types

To assess the ability of our aMIL approach to predict levels of the TGFβ-CAF signature in H&E-stained tissue images, we deployed the model on WSI from six TCGA datasets (BLCA, BRCA, STAD, LUAD, LUSC, PRAD). Model outputs consisted of a slide-level continuous TGFβ-CAF score (Fig. 1B). aMIL models were evaluated based on their continuous slide-level TGFβ-CAF scores in comparison to computed TGFβ-CAF scores from transcriptomic data using Spearman ⍴ values (Figure 2A, Supplementary Figure S8A), receiver operating characteristic curves (Figure 2B, Table 1) and dichotomized slide-level predictions (Table 1). The performance of these aMIL models varied by indication, with BLCA, BRCA, STAD and PRAD models exhibiting good performance based on continuous slide-level score (AUROC ≥ 0.75), while the models specific to LUAD and LUSC demonstrated comparatively weaker performance (AUROC < 0.65). The resulting confusion matrices are shown in Figure 2C and Supplementary Figure S8C.

**Figure 2.**
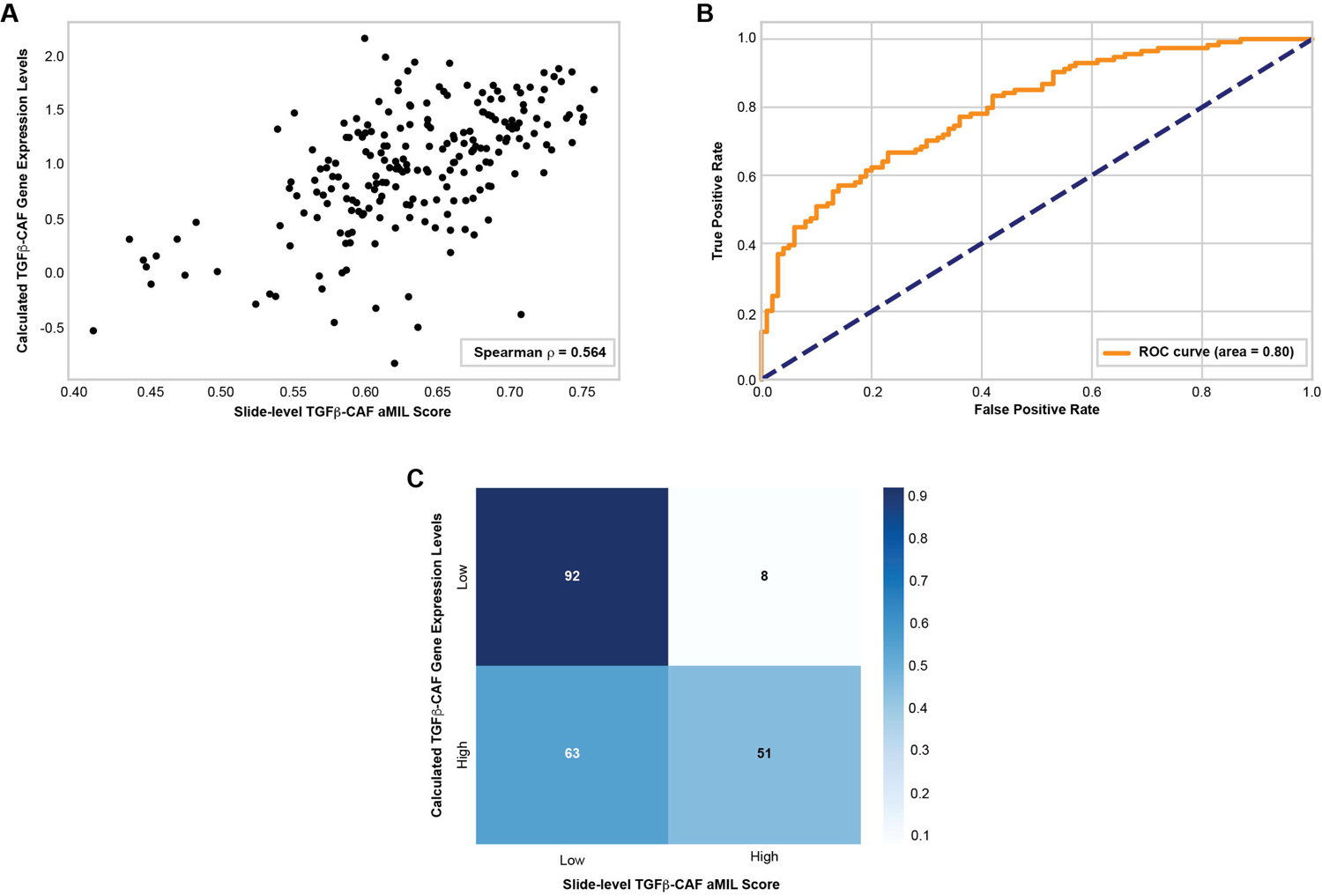
Model performance in predicting TGFβ-CAF levels in breast cancer. A) Correlation of model-predicted slide-level TGFβ-CAF scores with levels calculated from sequencing data for each case (Spearman r = 0.567). B) Performance of aMIL model in predicting TGFβ-CAF levels in BRCA (AUROC = 0.80). C) Confusion matrix depicting agreement of model predictions with ground truth. The number of slides in each category is indicated. All analyses were performed in the test set.

**Table 1.** Model performance in test set for predicting TGFβ-CAF in six cancer subtypes.

|  | **BLCA** | **BRCA** | **STAD** | **LUAD** | **LUSC** | **PRAD** |
| --- | --- | --- | --- | --- | --- | --- |
| **Number of Slides** |  |  |  |  |  |  |
| **Training** | 277 | 596 | 249 | 316 | 269 | 591 |
| **Validation** | 93 | 185 | 87 | 104 | 93 | 105 |
| **Test** | 85 | 214 | 81 | 107 | 91 | 118 |
| **AUROC (Validation set)** | 0.78 | 0.78 | 0.74 | 0.77 | 0.67 | 0.76 |
| **AUROC (Test set)** | 0.75 | 0.80 | 0.82 | 0.64 | 0.62 | 0.75 |
| **Accuracy** | 0.68 | 0.67 | 0.73 | 0.66 | 0.52 | 0.65 |
| **Mis-Classification** | 0.32 | 0.33 | 0.27 | 0.34 | 0.48 | 0.35 |
| **Sensitivity** | 0.61 | 0.45 | 0.75 | 0.87 | 0.58 | 0.67 |
| **Specificity** | 0.74 | 0.92 | 0.71 | 0.31 | 0.48 | 0.65 |
| **Precision** | 0.66 | 0.86 | 0.68 | 0.69 | 0.39 | 0.48 |
| **F1 Score** | 0.63 | 0.59 | 0.71 | 0.77 | 0.46 | 0.56 |

These models were also compared against negative control models to quantify the signal gained by training on these TGFβ-CAF labels. A model trained on slide background patches yielded AUROC=0.53 (Figure S9A), while five models trained with randomly permuted ground truth labels resulted in AUROC values between 0.45-0.53 (Figure S9B). These negative control model values are significantly lower than those derived from the cancer-specific aMIL models described (p ≤ 0.001, DeLong’s test), increasing our confidence that model performance is not driven by confounding signals present in imaging data.

### Features associated with TGFβ-CAF model attention in BRCA

To investigate the underlying morphological and spatial patterns driving TGFβ-CAF predictions, we performed in-depth analysis on the BRCA dataset, given the strong performance of this aMIL model. To investigate the cell and tissue features that drive model attention, we employed the strategy outlined in Figure 1B. In slides predicted to be TGFβ-CAF-high, features were extracted from regions for which the aMIL score was in the top 25th percentile (“excitatory”); likewise, in slides predicted to be TGFβ-CAF-low, features were extracted from regions for which the aMIL score was in the bottom 25th percentile (“inhibitory”). Features derived from these excitatory and inhibitory regions were then directly compared.

In addition to the aMIL TGFβ-CAF model, PathExplore models, which provide pixel-level predictions of tissue regions (Supplementary Figure 4) and cell types^18^ in cancer WSI, were deployed on WSI from the TCGA BRCA cohort (Figure 3A). To determine the cell and tissue features associated with TGFβ-CAF model attention, we extracted the features (N=297) from aggregated excitatory regions in slides predicted to be TGFβ-CAF high and from aggregated inhibitory regions in slides predicted to be TGFβ-CAF low. Clear separation between model-predicted TGFβ-CAF-high and -low was observed when performing dimensionality reduction on these cell and tissue features of high-contribution regions, with 55% of variation explained by the first two principal components (Figure 3B). In particular, density of immune cells in cancer stroma was the most important feature driving PC1, while the relative proportion of cancer epithelial cells in the tumor and the relative area of cancer stroma to total tissue were the most important features driving PC2. Many features, most relating to fibroblasts or immune cells, were observed to significantly differ according to TGFβ-CAF status (Table 2).

**Figure 3.**
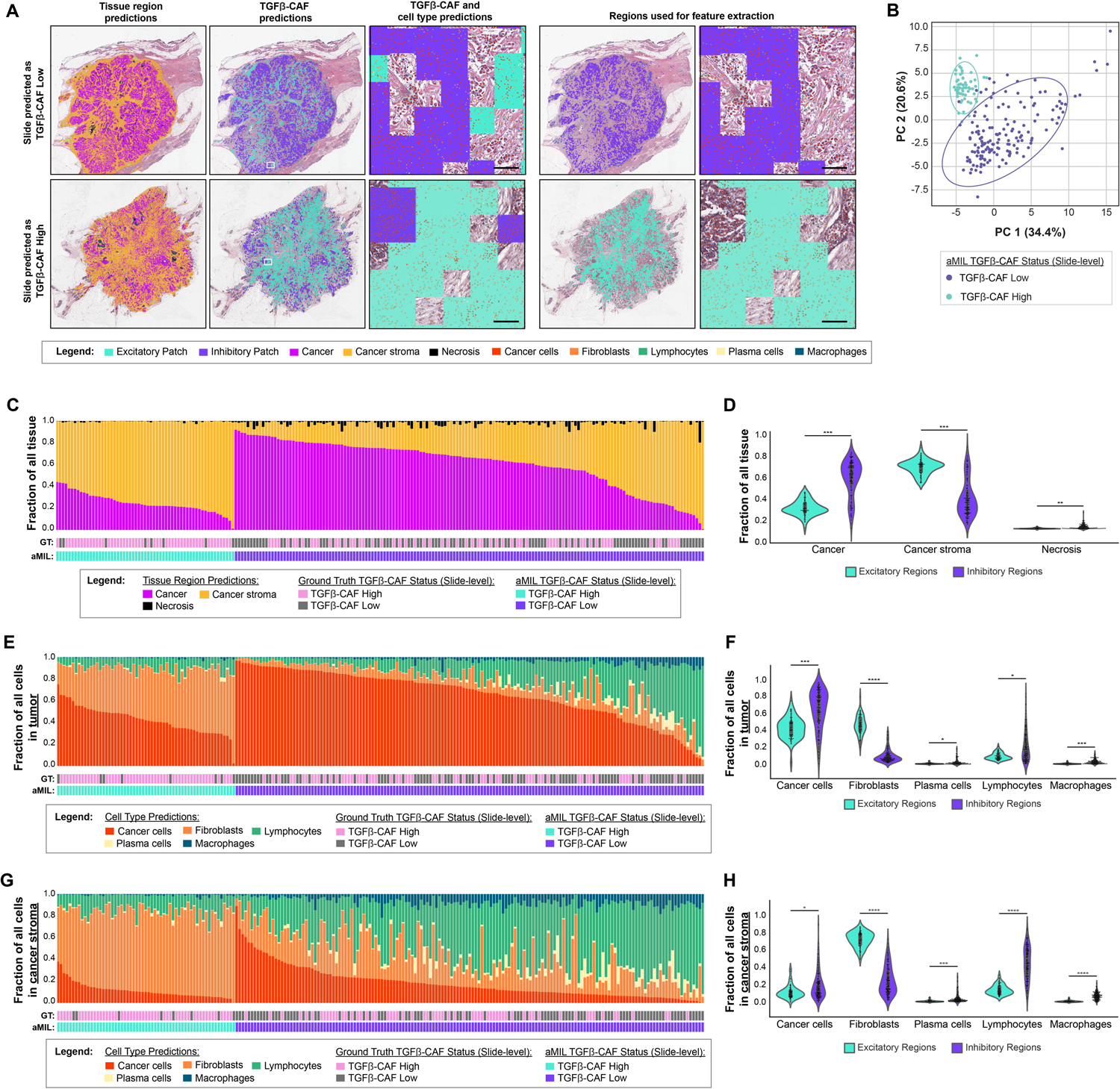
Quantification of tissue regions and cell types between excitatory regions in slides predicted to be TGFβ-CAF-high and inhibitory regions in slides predicted to be TGFβ-CAF-low. A) Overlays in BRCA WSI depicting predictions of tissue regions, cell types, and TGFβ-CAF levels in slides predicted to be TGFβ-CAF-high and TGFβ-CAF-low. Features were extracted from excitatory regions in TGFβ-CAF-high slides and inhibitory regions in TGFβ-CAF-low slides. High magnification images were captured from the regions indicated by insets. Scale bars indicate 0.1 mm. B) Principal components analysis using HIFs extracted from high contribution patches. A projection of patches onto the first two principal components, colored by predicted TGFβ-CAF status, is shown. C) Proportional areas of cancer epithelium, cancer-associated stroma, and necrosis in TGFβ-CAF high and TGFβ-CAF low slides. TGFβ-CAF status, both ground truth (GT) and aMIL-predicted, are shown. D) Quantification of relative proportional areas of tissue regions in slides predicted to be TGFβ-CAF high and TGFβ-CAF low. E) Relative counts of cancer epithelial cells, fibroblasts, plasma cells, lymphocytes, and macrophages in tumor regions of TGFβ-CAF high and TGFβ-CAF low slides. F) Quantification of relative cell counts in tumor regions of TGFβ-CAF high and TGFβ-CAF low slides. G) Relative counts of cancer epithelial cells, fibroblasts, plasma cells, lymphocytes, and macrophages in stromal regions of TGFβ-CAF high and TGFβ-CAF low slides. H) Quantification of relative cell counts in tumor regions of TGFβ-CAF high and TGFβ-CAF low slides. Asterisks in panels indicate FDR-adjusted p-values calculated by Mann Whitney U test, as follows: ****: p<1e-20; ***: p< 1e-10; **: p< 1e-5; *: p< 0.05.

**Table 2.** Comparison of TME HIFs and Nuclear HIFs in high contribution regions between TGFβ-CAF high and low predicted slides. The top ten cell and tissue features and nuclear features are shown. The AUC values shown quantify the effect size for the Mann-Whitney U test.

|  | Feature | AUC | P value | Adjusted p value | Predicted TGFβ-CAF status of regions enriched for feature |
| --- | --- | --- | --- | --- | --- |
| **Cell and Tissue Features** | Fraction of fibroblasts relative to all predicted cells in the tumor | 0.992 | $4.77 \times 10^{-29}$ | $1.81 \times 10^{-27}$ | High |
| | Fraction of fibroblasts relative to all predicted cells in the cancer stroma | 0.991 | $7.02 \times 10^{-29}$ | $1.81 \times 10^{-27}$ | High |
| | Density of fibroblasts in the tumor | 0.990 | $1.00 \times 10^{-28}$ | $1.81 \times 10^{-27}$ | High |
| | Density of macrophages in the cancer stroma | 0.955 | $4.35 \times 10^{-25}$ | $5.88 \times 10^{-24}$ | Low |
| | Fraction of macrophages relative to all predicted cells in the cancer stroma | 0.949 | $1.77 \times 10^{-24}$ | $1.91 \times 10^{-23}$ | Low |
| | Fraction of immune cells relative to all predicted cells in the cancer stroma | 0.947 | $3.08 \times 10^{-24}$ | $2.77 \times 10^{-23}$ | Low |
| | Density of fibroblasts in the cancer epithelium | 0.939 | $1.68 \times 10^{-23}$ | $1.22 \times 10^{-22}$ | High |
| | Fraction of lymphocytes relative to all predicted cells in the cancer stroma | 0.939 | $1.81 \times 10^{-23}$ | $1.22 \times 10^{-22}$ | Low |
| | Density of fibroblasts in the cancer stroma | 0.936 | $3.52 \times 10^{-23}$ | $2.11 \times 10^{-22}$ | High |
| | Fraction of fibroblasts relative to all predicted cells in the cancer epithelium | 0.925 | $4.39 \times 10^{-22}$ | $2.37 \times 10^{-21}$ | High |
| **Nuclear Features** | Mean lymphocyte nucleus eccentricity | 0.981 | $1.91 \times 10^{-23}$ | $4.21 \times 10^{-21}$ | High |
| | Standard deviation of lymphocyte nucleus eccentricity | 0.978 | $3.66 \times 10^{-23}$ | $4.21 \times 10^{-21}$ | Low |
| | Standard deviation of fibroblast nucleus perimeter length | 0.937 | $9.81 \times 10^{-20}$ | $7.52 \times 10^{-18}$ | High |
| | Standard deviation of fibroblast nucleus major axis length | 0.925 | $9.75 \times 10^{-19}$ | $5.60 \times 10^{-17}$ | High |
| | Mean lymphocyte nucleus circularity | 0.900 | $8.59 \times 10^{-17}$ | $3.95 \times 10^{-15}$ | Low |
| | Standard deviation of fibroblast nucleus area | 0.896 | $1.52 \times 10^{-16}$ | $5.53 \times 10^{-15}$ | High |
| | Standard deviation of fibroblast nucleus minor axis length | 0.896 | $1.68 \times 10^{-16}$ | $5.53 \times 10^{-15}$ | High |
| | Standard deviation of fibroblast nucleus circularity | 0.832 | $3.80 \times 10^{-12}$ | $1.90 \times 10^{-10}$ | High |
| | Standard deviation of lymphocyte nucleus perimeter length | 0.779 | $4.71 \times 10^{-9}$ | $1.20 \times 10^{-7}$ | High |
| | Mean fibroblast nucleus minimum grayscale intensity value | 0.764 | $2.77 \times 10^{-8}$ | $6.38 \times 10^{-7}$ | High |

These results suggest that spatial variation in tissue regions and cell types may explain aMIL TGFβ-CAF model attention. Interestingly, visual examination of the PathExplore and aMIL heatmaps revealed apparent co-localization of excitatory regions with cancer stroma and inhibitory regions with cancer epithelium (Figure 3A). In addition, cancer cells appeared to be enriched in inhibitory regions, and fibroblasts appeared to be enriched in excitatory regions. Therefore, we sought to better understand the relative proportions of tissue regions and cell types in regions determined to be excitatory or inhibitory. To do so, PathExplore features quantifying the relative areas of predicted tissue regions were extracted and compared between excitatory regions of model-predicted TGFβ-CAF-high slide and inhibitory regions of model-predicted TGFβ-CAF-low slides. Indeed, as shown in Figures 3C and 3D, excitatory regions showed increased area of cancer stroma relative to total tissue (p<0.001), while inhibitory regions showed increased area of cancer epithelium relative to total tissue (p<0.001). Increased necrosis was also observed in inhibitory regions (p<0.001). Across tumor tissue (e.g., combined cancer and stroma areas), cancer cells (p<0.001), plasma cells (p<0.05), lymphocytes (p<0.05), and macrophages (p<0.001) were enriched in inhibitory regions, while fibroblasts were enriched in excitatory regions (p<0.001) (Figure 3E-F). When restricting these analyses to the stroma, we observed similar trends. Immune cells – plasma cells (p<0.001), lymphocytes (p<0.001), and macrophages (p<0.001) – were particularly enriched in inhibitory regions, while fibroblasts were enriched in excitatory regions (p<0.001) (Figure G-H).

In addition, we aimed to determine the extent to which cell-level nuclear morphology corresponds with TGFβ-CAF model predictions in breast cancer. To do so, we extracted and compared cell type-specific nuclear features in excitatory and inhibitory regions. Interestingly, features relating to lymphocyte nuclear shape, particularly eccentricity and circularity, were observed to differ between excitatory and inhibitory regions (Figure 4, Table 2). Furthermore, features quantifying fibroblast nuclear size, including standard deviation of nucleus perimeter, standard deviation of nucleus area, standard deviation of nucleus major axis length, and standard deviation of nucleus minor axis length, differed between excitatory and inhibitory regions (Figure 4, Table 2).

**Figure 4.**
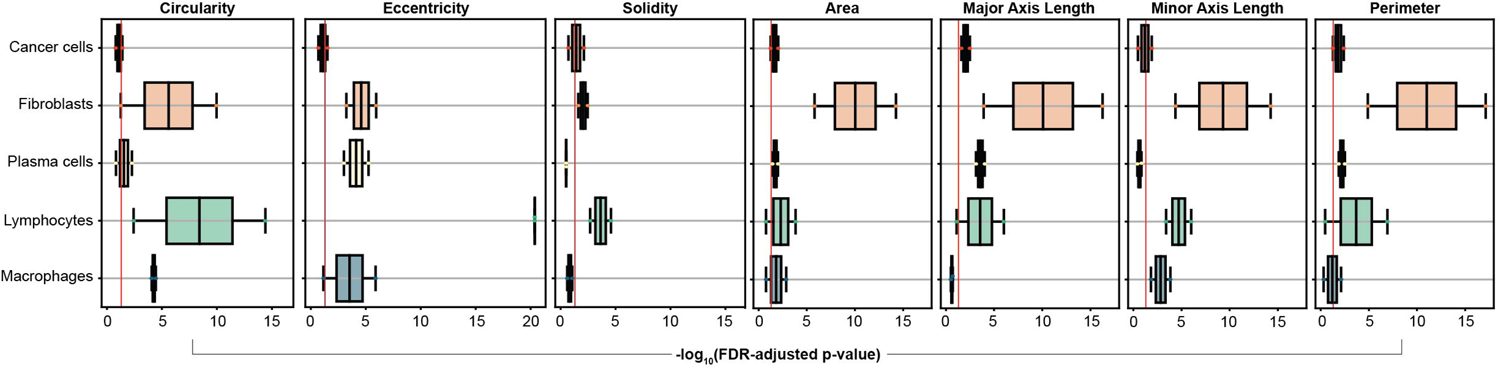
Association of nuclear morphology with TGFβ-CAF model attention in BRCA. Quantification of cell type-specific nuclear features associated with excitatory and inhibitory regions. Boxes represent the interquartile range (IQR) of -log10 FDR adjusted p-values, with the line inside box indicating the median, and whiskers extend to the smallest and largest data points within 1.5*IQR from the first and third quartiles, respectively.

We also sought to determine whether performing these analyses at the whole-slide level recapitulates the results obtained from the focusing on regions of interest. Notably, TME features extracted from high-contribution regions show a more pronounced difference than those extracted from whole slides (Supplementary Figure S10A-C). Similarly, nuclear features extracted from high contribution regions show a more pronounced difference than those extracted from whole slides (Supplementary Figure S10D,E). These differences demonstrate the ability of aMIL to reveal novel morphological correlates of gene expression patterns, which traditional quantitative tissue analysis cannot identify.

### Contribution of immune infiltration to TGFβ-CAF predictions in breast cancer

LRRC15+ CAFs have previously been shown to be associated with extracellular matrix remodeling and collagen formation in breast cancer^10^. Given this prior observation and the differences in fibroblast nuclear morphology that were associated with TGFβ-CAF predictions, we hypothesized that the organization of the tumor stroma may also correspond with model attention. To address this question, models trained to predict the stromal composition and organization from an H&E-stained WSI^21^ were applied on slides in the dataset. The stromal subtypes predicted by this model were: densely inflamed stroma (characterized by high degrees of immune cell infiltration), mature stroma (characterized by the presence of mature, elongated collagen fibers^22^), immature stroma (characterized by a lack of mature collagen fibers), fibroblastic stroma (characterized by high degrees of fibroblast infiltration) and elastosis (characterized by the presence of elastin fibers) (Figure 5A). Consistent with our previous results regarding lymphocytes and TGFβ-CAF model attention, inhibitory regions had higher proportions of densely inflamed stroma; in contrast, excitatory regions had more mature stroma, fibroblastic stroma, and elastosis (Figure 5B). To better understand the relationship between predicted TGFβ-CAF and stromal subtypes, we assessed the cell types present within each subtype in excitatory and inhibitory regions (Figure 5C-G). Within each subtype, immune cells (especially lymphocytes) were enriched in inhibitory compared to excitatory regions; conversely, fibroblasts were enriched in excitatory regions compared to inhibitory regions.

**Figure 5.**
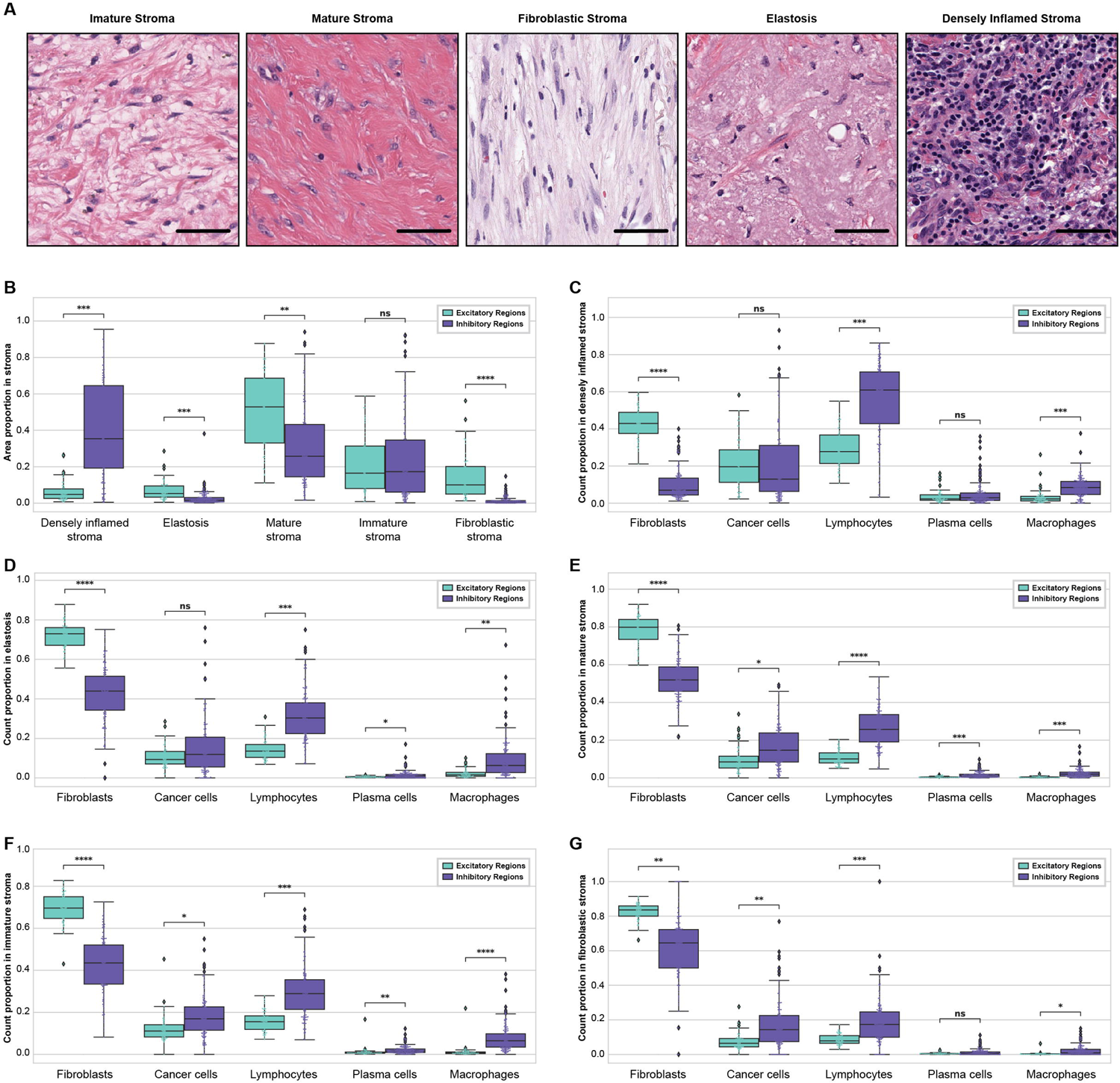
Association of stromal features extracted from high-contribution regions with predicted TGFβ-CAF levels in BRCA. A) Representative example images from H&E-stained WSI of the five predicted stromal subtypes. B) Quantification of the relative area of each predicted stromal subtype in slides predicted to be TGFβ-CAF high or low. Relative numbers of each predicted cell type are shown for C) densely inflamed stroma, D) elastosis, E) mature stroma, F) immature stroma, and G) fibroblastic stroma. Asterisks in panels C, E, and G indicate FDR-adjusted p-values calculated by Mann Whitney U test, as follows: ****: p<1e-20; ***: p< 1e-10; **: p< 1e-5; *: p< 0.05.

Previous research into LRRC15+ fibroblasts and the TGFβ-CAF gene signature revealed an association with immunosuppressive activity and poor response to immunotherapy^8,9^. Our results appear to support these findings, as excitatory regions harbored significantly fewer lymphocytes in both the stroma and whole tumor (Figure 3) and were significantly reduced in stroma predicted to be densely inflamed (Figure 5). To further address the relationship between immunophenotype and TGFβ-CAF model attention, we assessed the cell types present within close proximity (120 μm) of the cancer epithelium-stroma interface (ESI). On both the cancer (Supplemental Figure S11A) and stromal (Supplemental Figure S11B) sides of the ESI, both cancer and immune cells were enriched in inhibitory regions, while fibroblasts were enriched in excitatory regions. These results further suggest that high levels of TGFβ-CAF may prevent lymphocytes from approaching and infiltrating into the cancer epithelium.

### Validation of TGFβ-CAF morphologic correlates in a pan-cancer analysis

Having shown that aMIL allows the identification of tumor features that correspond with TGFβ-CAF predictions in breast cancer, we sought to determine if similar cell and tissue features contribute to TGFβ-CAF prediction in additional cancer types. PathExplore models were additionally deployed on WSI from the BLCA, STAD, LUAD, LUSC, and PRAD TCGA datasets, as the TGFβ-CAF signature has potential relevance in these cancer subtypes^8^. As with analysis of the BRCA dataset alone, these results revealed the enrichment of cancer stroma in excitatory regions and enrichment of cancer epithelium in inhibitory regions (Figure 6A). Features quantifying the relative areas of predicted tissue regions were extracted and compared between excitatory and inhibitory regions, revealing associations between the relative amounts of cancer stroma and cancer epithelium with high and low TGFβ-CAF model attention, respectively (Figure 6B). Further exploration of the extracted cell type features revealed enrichment of cancer epithelial cells and lymphocytes in inhibitory regions and enrichment of fibroblasts in excitatory regions (Figure 6C, D). Similarly, in stromal tissue, fibroblasts were enriched in excitatory regions and lymphocytes were enriched in inhibitory regions (Figure 6E, F).

**Figure 6.**
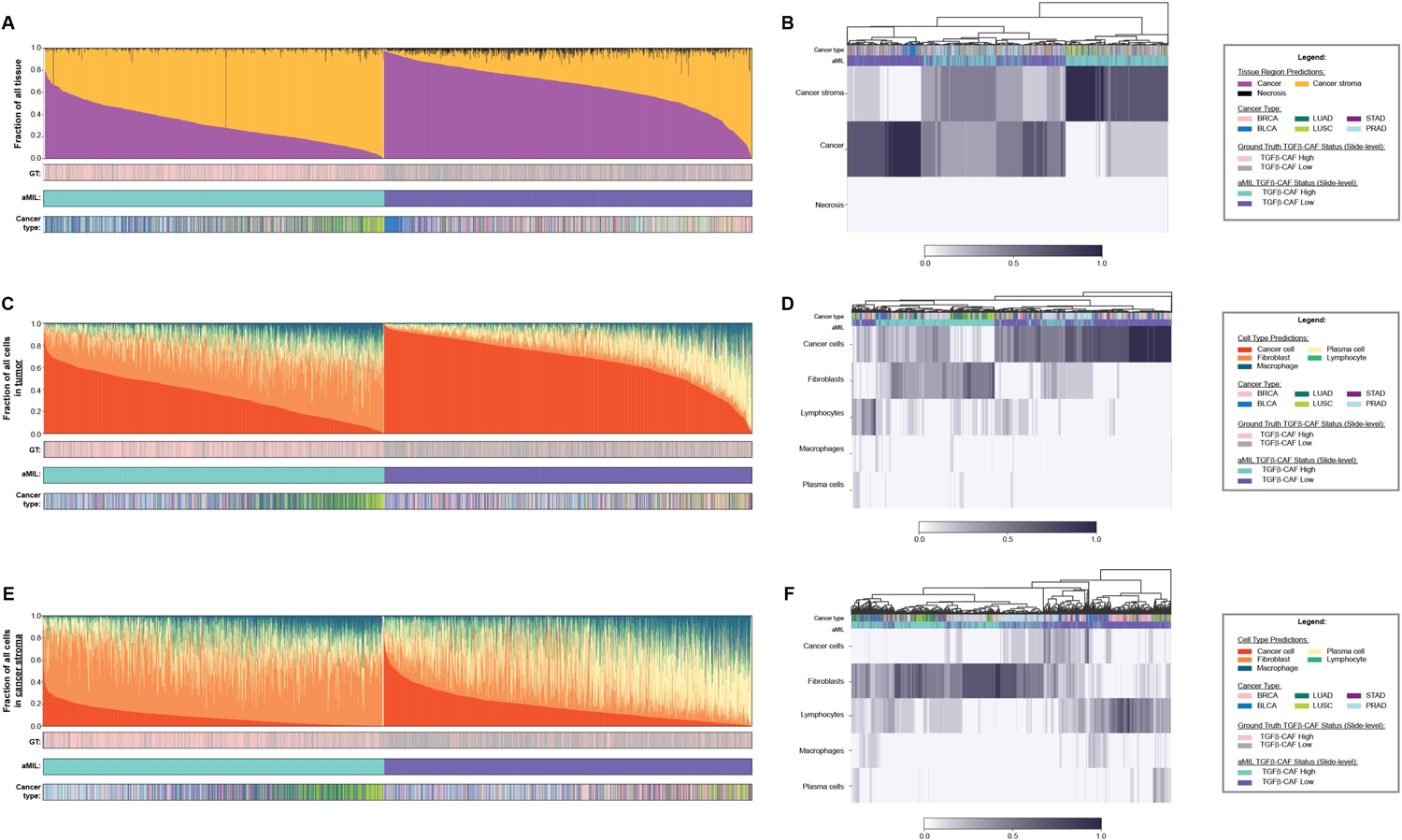
Quantification of tissue regions and cell types found in TGFβ-CAF-high and TGFβ-CAF-low regions across cancer types. A) Proportional areas of cancer epithelium, cancer-associated stroma, and necrosis in TGFβ-CAF high and TGFβ-CAF low slides, with hierarchical clustering (B). TGFβ-CAF status, both ground truth (GT) and aMIL-predicted, are shown. C) Relative counts of cancer epithelial cells, fibroblasts, plasma cells, lymphocytes, and macrophages in tumor regions of TGFβ-CAF high and TGFβ-CAF low slides, with hierarchical clustering (D). E) Relative counts of cancer epithelial cells, fibroblasts, plasma cells, lymphocytes, and macrophages in stromal regions of TGFβ-CAF high and TGFβ-CAF low slides, with hierarchical clustering (F).

All in all, we observed several consistent patterns across the tumor types examined. Across bladder, breast, stomach, lung, and prostate cancers, a higher relative area of cancer tissue and enrichment of cancer epithelial cells was observed in regions contributing most to TGFβ-CAF-low predictions. Furthermore, in all cancer types examined, a higher relative area of stromal tissue and enrichment of fibroblasts in both tumor and stroma was observed in regions contributing most to TGFβ-CAF-high predictions. Fibroblast nuclear features were shown to differ between these regions in all cancer types, while lymphocyte nuclear features differed between these regions in the BRCA, LUAD, and LUSC datasets. These results, summarized in Table 3, indicate that similar cell and tissue characteristics contribute to TGFβ-CAF prediction across cancer types. Given that the aMIL model works across indications, these results suggest that the underlying biology of regions expressing TGFβ-CAF is similar in multiple solid tumor subtypes.

**Table 3.** Summary of associations observed between features and TGFβ-CAF predictions across cancer types.

| Observation | Indication(s) |
| --- | --- |
| Higher relative area of Stroma in Excitatory regions | ALL |
| Higher relative area of Cancer in Inhibitory regions | ALL |
| Higher relative number of Fibroblasts in Tumor/Stroma in Excitatory regions | ALL |
| Higher relative number of Cancer Epithelial Cells in Tumor in Inhibitory regions | ALL |
| Difference in Fibroblast nuclear size features between Excitatory and Inhibitory regions | ALL |
| Difference in Lymphocyte nuclear shape features between Excitatory and Inhibitory regions | BRCA, LUAD, LUSC |

## Discussion

Molecular subtyping of CAFs has yielded a wealth of information regarding the impact of these cells on tumor progression and response to therapy. However, the identification of clinically relevant CAF subtypes (e.g., LRCC15+ CAFs) necessitates transcriptomic analyses of tumor samples at the single-cell level, hindering the adoption of these subtypes as clinical biomarkers. Here, we describe a novel approach, aMIL, which allows the interpretable association between histopathology and transcriptional signatures, resulting in spatially localized model-predicted GES.

The aMIL approach that we describe herein is accurate for predicting levels of the TGFβ-CAF GES from bulk RNA sequencing using H&E-stained WSI in multiple cancer types. Moreover, by deploying additional ML models on the same WSI to predict tissue regions, cell types, and stromal subtypes, we unlock avenues for additional experimentation, interpretation and further investigation of underlying biology. Importantly, this approach provides us with the ability to look inside the proverbial “black box” to understand the cell and tissue features with the greatest association with model predictions. Tissue regions contributing most to TGFβ-CAF-high predictions (“excitatory”) were preferentially localized to cancer stroma regions with high fibroblast density. In contrast, tissue regions contributing most to TGFβ-CAF-low predictions (“inhibitory”) were preferentially localized to cancer epithelium and stromal regions with high lymphocyte infiltration. Furthermore, TGFβ-CAF excitatory regions were associated with distinct stromal subtypes (fibroblastic stroma and mature stroma), while TGFβ-CAF inhibitory regions were prevalent in densely inflamed stroma.

Notably, as summarized in Figure 7, the features that we observe to associate most with model predictions match well with previous observations of the LRRC15+ fibroblasts and the associated TGFβ-CAF and ecm-myCAF gene signatures. First, these gene signatures were derived from and characterize cancer-associated fibroblasts^8^, and we observe excitatory regions to be localized to areas of cancer-associated stroma with abundant fibroblasts. These results suggest that our model is focused on the appropriate cell types for the prediction of high levels of the TGFβ-CAF gene signature. *LRRC15* is a component of the ecm-myCAF gene signature in breast cancer, which is characterized by increased collagen synthesis^10^. Interestingly, the TGFβ-CAF gene signature, itself, contains many genes involved in ECM remodeling and collagen deposition and/or arrangement, including *MMP11*, *COL11A1*, *CTHRC1*, and *COL12A1*^8^. We observe that tissue regions with elevated contribution to prediction of TGFβ-CAF-high have high amounts of stroma characterized by the presence of mature, elongated collagen fibers. This result supports the idea that our aMIL model attention is focused on regions enriched for ecm-myCAF population, known to be associated with LRRC15+ fibroblasts. Finally, prior work has demonstrated that CAFs expressing LRRC15 are associated with immunosuppressive activity and poor response to immunotherapy^7–10^. We observed that inhibitory regions contained a high proportion of immune cells, particularly lymphocytes, and were localized to regions of densely inflamed cancer stroma.

**Figure 7.**
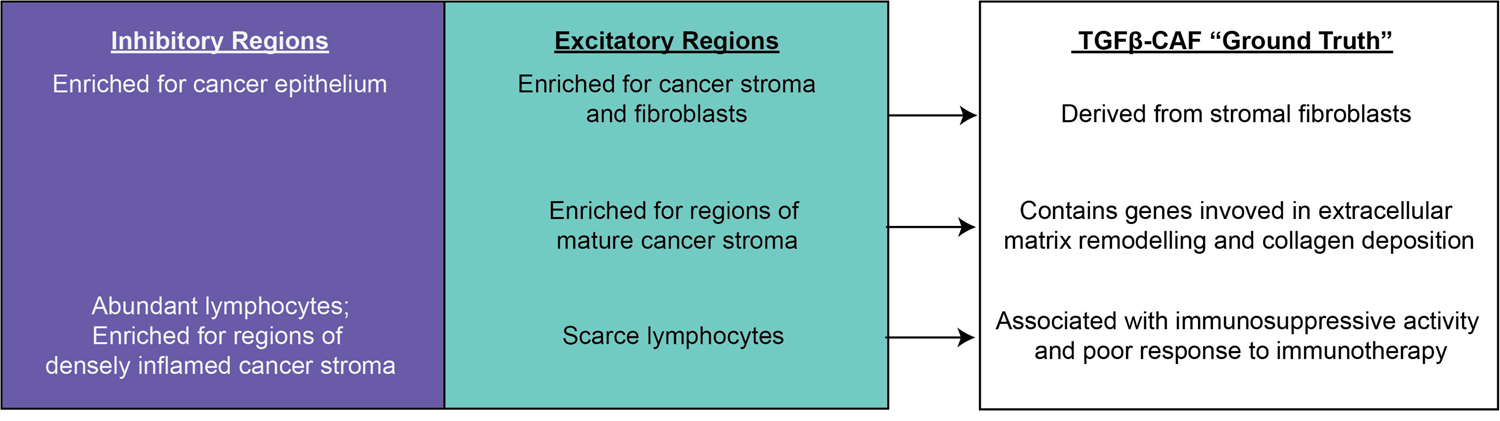
Summary of features associated with inhibitory and excitatory patches and relationship with known TGFβ-CAF characteristics.

That the features corresponding to our model’s attention recapitulate previously reported observations regarding TGFβ-CAF adds to our confidence that our model is focusing on biologically relevant regions of cancer tissue, allowing us to further speculate about the biological and translational relevance of our findings. For instance, our observation that excitatory regions were associated with mature collagen and a lower density of lymphocytes is intriguing in light of the potential link between ECM composition and anti-tumor immunity. Indeed, collagen fiber orientation within the tumor microenvironment has previously been shown to directly impact the degree of lymphocyte infiltration into the cancer epithelium^23–25^. Furthermore, a positive correlation has been described between ecm-myCAFs and infiltration of PD-1+, CTLA4+, and TIGIT+ CD4+ T lymphocytes, as well as FOXP3+ regulatory T cells and a negative correlation with CD8+ T lymphocytes^10^. Thus, ecm-myCAFs are associated with an immunosuppressive tumor microenvironment. While our models cannot discriminate between these lymphocyte subtypes from an H&E image, we do observe characteristics of an immunosuppressive microenvironment in regions found to contribute most to model predictions in slides predicted to be TGFβ-CAF-high. These excitatory regions show a dearth of lymphocytes (compared to inhibitory) and are negatively associated with stromal regions predicted to be densely inflamed. Notably, we observed distinct enrichment of nuclear features between lymphocytes in excitatory and inhibitory regions, potentially reflecting distinct lymphocyte subpopulations (e.g., inhibitory Tregs, PD-1 positive cells in excitatory regions, etc.). Therefore, our results seemingly confirm the difference in lymphocyte composition that results from disparate TGFβ-CAF levels and suggest that nuclear morphology may differ between these cells (“form follows function”). However, further work is needed to better understand the morphologic differences between lymphocyte subtypes in this regard.

Interestingly, in the BRCA TCGA dataset, features associated with fibroblast nuclear size were previously associated with GES that mirror the results observed herein^18^. Larger fibroblast nuclei were positively associated with pathways involved in ECM remodeling and collagen organization and were negatively associated with pathways involved in anti-tumor immunity. In this study, we noted a similar association between fibroblast nuclear features and TGFβ-CAF predictions in BRCA: features quantifying fibroblast nuclear size differed between excitatory and inhibitory regions. We also observed these regions to differ in their stromal subtype and immune infiltration patterns, such that excitatory regions were more likely to display mature collagen deposition and had a relative scarcity of lymphocytes. Therefore, the results presented herein confirm and extend our previous observations regarding fibroblast nuclear morphology in breast cancer. Furthermore, these results are consistent with prior observations of ecm-myCAFs, which are known to express LRRC15 and are associated with ECM remodeling, collagen organization, and poor immune infiltration^10^.

Given the relationship between LRRC15+ CAFs and immunotherapy response, there have been efforts to target this CAF population; notably, an antibody-drug conjugate targeting LRRC15, ABBV-085, is already in clinical trials^26,27^. A recent clinical trial assessed safety and preliminary antitumor efficacy of this agent in patients with multiple tumor types^26^. The most prevalent cancer type was sarcoma, which expresses LRRC15 given the mesenchymal nature of the tumor cells, but a small number of patients with epithelial tumors (including breast cancer) were also enrolled in the trial. While responses were rare in epithelial tumors, evidence of increased T cell infiltration was observed in non-responding patients with breast cancer treated with ABBV-085, suggesting that the agent is, indeed, targeting LRRC15 fibroblasts, albeit without an observable clinical benefit. Interestingly, there was no clear correlation between LRRC15 expression and response. The authors suggested that assay sensitivity and tumor heterogeneity may have contributed to this result. The approach that we describe herein may have potential to better understand the link between LRRC15+ expression, using predicted TGFβ-CAF levels in H&E-stained tissue as a surrogate, and response to LRRC15 inhibition.

Despite the advantages of our approach, this study is not without limitations. Most notably, to assess the accuracy of our aMIL TGFβ-CAF predictions, we were limited by cohorts of publicly available H&E-stained WSI with associated transcriptomic data. Thus, we lack a true out-of-domain dataset in this regard, since our cohorts were entirely derived from the TCGA database. That said, we observed high model accuracy in multiple tumor types, which bodes well for the generalizability of our approach. Furthermore, while the aMIL approach performed well in BLCA, BRCA, STAD and PRAD (AUROC ≥ 0.75), the aMIL models specific to LUAD and LUSC struggled comparatively (AUROC < 0.65). Results from multiple negative control models further suggest that our highest performing aMIL models are able to detect true biological TGFβ-CAF signal in these slides. Interestingly, the fact that the lowest performing TGFβ-CAF aMIL model (LUSC, AUROC=0.62) significantly outperformed the negative control models suggests that it is also detecting some degree of underlying signal not present in the negative control labels. The interpretability approach used here can also be applied to troubleshooting poor model performance, by flagging features or areas on which the models are inappropriately focused. While we focused our in-depth analysis on results from a single high-performing model (BRCA), this approach could be similarly applied to a low-performing model to identify features that are informing its predictions, potentially identifying confounders. We also report interpretability results for all six aMIL models in a pan-cancer manner. Interestingly, despite the inclusion of the sub-optimal LUAD and LUSC models in the pan-cancer analysis, the features identified as contributing most to the prediction of TGFβ-CAF-high and -low were similar to those identified in BRCA.

While other models have been described previously that predict GES levels from H&E-stained images^28–31^, the aMIL approach described in this study stands out for several reasons. First, the aMIL model can generate spatially resolved predictions of labels (e.g., gene expression levels) and therefore yield imputed transcriptomic predictions at both the slide level as well in more focused regions, enabling a more comprehensive understanding of the within-tumor heterogeneity of gene expression patterns and the effects of local enrichment. Moreover, the integration of additional slide-based analyses, such as models predicting tissue regions, cell types, and stromal subtypes, enhances the interpretability of aMIL predictions. By combining these data-driven approaches, previously considered “black-box” models become interpretable in ways that hold clinical relevance. This holistic approach has the potential to not only elucidate the underlying mechanisms driving gene expression patterns but also facilitate the translation of these findings into clinically actionable insights.

Therefore, aMIL, a novel ML framework, has the ability not only to predict gene expression in a spatially localized manner in an H&E-stained image, but also to yield interpretability through the association of these expression patterns with localized histopathological features. Given the challenges of assessing gene expression for use as a clinical biomarker, the approach described herein may have potential for the assessment of transcriptional signatures using histopathology specimens.

## ACKNOWLEDGEMENTS

The authors would like to thank the software engineering and machine learning operations teams at PathAI for developing the systems and pipelines used for model development and feature extraction. The authors also thank GCI Health for assistance with figure design.

## FUNDING

This work was funded by PathAI, Inc.

## AUTHOR CONTRIBUTIONS

Conceptualization: MM, JK, ATW, CP

Methodology: MM, ZG, SAJ, DJ, HP, AK, CP

Investigation: MM, JK, ZG, YG

Visualization: MM, JK, JBC

Funding acquisition: N/A

Project administration: BR, JA, SH, AK, ATW, CP

Supervision: JA, AK, ATW, CP

Writing – original draft: MM, JK, JBC, CP

Writing – review & editing: MM, JK, ZG, YG, JBC, SAJ, DJ, HP, LY, BR, JA, SH, AK, ATW, CP

## DATA AND MATERIALS AVAILABILITY

Histopathology images from the Cancer Genome Atlas dataset are available at https://www.cancer.gov/about-nci/organization/ccg/research/structural-genomics/tcga. Images and annotations of nuclei from the following datasets will be made available on GitHub prior to publication: training set, validation set, test set. Access to feature tables, cell-, tissue-, and nuclei-type heatmaps, as well as usage of cell- and tissue-type classification models, are available upon reasonable request to academic investigators without relevant conflicts of interest for non-commercial use who agree not to distribute the data. Access requests can be made to.

## CODE AVAILABILITY

Model parameters for cell, tissue, aMIL, and stromal models, and codes for model training, inference, and feature extractions are not disclosed. Access requests for such a code will not be considered to safeguard PathAI’s intellectual property. All source code for reproducing statistical analyses and molecular predictions, will be deposited to GitHub prior to publication, and the link will be provided at that time.

## Notes

### Competing Interest Statement

MM, JK, ZG, YG, JBC, SAJ, DJ, HP, LY, BR, JA, SH, AK, ATW, and CP are current or former employees of and hold stock in PathAI, Inc. YG is a former employee of and holds stock in Finch Therapeutics.

