## Supplementary Table S1 for "Spatial mapping of immunosuppressive CAF gene signatures in H&E-stained images using additive multiple instance learning"

**Supplementary Table S1. Dataset characteristics**

| | BLCA (N=455) | BRCA (N=995) | ESCA & STAD (N=417) | LUAD (N=527) | LUSC (N=453) | PRAD (N=814) |
|---|---|---|---|---|---|---|
| **Pathologic stage, N (%)** | | | | | | |
| I | 3 (0.7) | 163 (16.4) | 53 (12.7) | 267 (50.7) | 225 (49.7) | |
| II | 155 (34.1) | 567 (57.0) | 128 (30.7) | 129 (24.5) | 130 (28.7) | |
| III | 148 (32.5) | 226 (22.7) | 185 (44.4) | 95 (18.0) | 88 (19.4) | |
| IV | 147 (32.3) | 17 (1.7) | 26 (6.2) | 27 (5.1) | 6 (1.3) | |
| Other | 2 (0.4) | 22 (2.2) | 25 (6.0) | 9 (1.7) | 4 (0.9) | 814 (100.0) |
| **Age at initial diagnosis, mean (SD)** | 67.9 (10.6) | 58.6 (13.3) | 64.7 (10.5) | 65.7 (10.2) | 67.7 (8.5) | 61.2 (7.0) |
| **Gender, N (%)** | | | | | | |
| Male | 335 (73.6) | 12 (1.2) | 292 (70.0) | 233 (44.2) | 341 (75.3) | 814 (100.0) |
| Female | 120 (26.4) | 983 (98.8) | 125 (30.0) | 294 (55.8) | 112 (24.7) | 0 (0.0) |
| **Survival status, N (%)** | | | | | | |
| 0 | 243 (53.4) | 856 (86.0) | 246 (59.0) | 315 (59.8) | 252 (55.6) | 787 (96.7) |
| 1 | 212 (46.6) | 139 (14.0) | 171 (41.0) | 212 (40.2) | 201 (44.4) | 27 (3.3) |
| **OS time (days), mean (SD)** | 819.2 (773.4) | 1262.8 (1215.8) | 612.8 (532.9) | 915.9 (850.5) | 1089.6 (1103.4) | 1184.7 (754.5) |
