## Supplementary Table S2 for "Spatial mapping of immunosuppressive CAF gene signatures in H&E-stained images using additive multiple instance learning"

**Supplementary Table 2. Median values used for binarization of TGFβ-CAF scores.**

| Indication | Threhsold |
|---|---|
| BLCA | 0.03782 |
| BRCA | 0.96693 |
| Gastric | 0.28861 |
| LUAD | 0.27165 |
| LUSC | 0.44829 |
| PRAD | -0.47574 |
