## Supplementary Table S3 for "Spatial mapping of immunosuppressive CAF gene signatures in H&E-stained images using additive multiple instance learning"

Supplementary Table 3. Summary of agreement (ICC) between model-derived cell counts and pathologists' cell counts.

| | BLCA (n=301 frames) | | | STAD (n=268 frames) | | |
|---|---|---|---|---|---|---|
| | Number of Cells | Model vs. Consensus [95% CI] | ΔICC [95% CI] | Number of Cells | Model vs. Consensus [95% CI] | ΔICC [95% CI] |
| **Cancer Cells** | 1548 | 0.67 [0.56, 0.76] | 0.1 [0.0, 0.13] | 2842 | 0.88 [0.86, 0.91] | -0.01 [-0.08, 0.06] |
| **Fibroblasts** | 615 | 0.62 [0.51, 0.73] | 0.07 [-0.02, 0.09] | 1666 | 0.5 [0.11, 0.7] | 0 [-0.08, 0.06] |
| **Lymphocytes** | 1962 | 0.79 [0.61, 0.88] | 0.04 [0.0, 0.09] | 3491 | 0.94 [0.93, 0.96] | 0.06 [0.03, 0.09] |
| **Macrophages** | 414 | 0.26 [0.06, 0.44] | 0.1 [0.03, 0.1] | 1210 | 0.03 [0.0, 0.11] | 0.01 [-0.04, 0.09] |
| **Plasma Cells** | 438 | -/68 [0.44. 0.81] | 0.06 [-0.04, 0.09] | 636 | 0.83 [0.79, 0.87] | 0.04 [-0.07, 0.10] |
