## Supplementary Table S4 for "Spatial mapping of immunosuppressive CAF gene signatures in H&E-stained images using additive multiple instance learning"

**Supplementary Table 4. Summar table of cell model performance characteristics.**

| | BLCA | | | | | |
|---|---|---|---|---|---|---|
| | **Recall** | | **Precision** | | **F1 Score** | |
| **Cell Class** | **Model vs. Avg Pathologist [95% CI]** | **Δrecall [95% CI]** | **Model vs. Avg Pathologist [95% CI]** | **ΔPrecision [95% CI]** | **Model vs. Avg Pathologist [95% CI]** | **ΔF1 [95% CI]** |
| **Cancer Cells (N = 1,548 cells)** | 0.62 [0.51, 0.68] | 0.05 [-0.03, 0.12] | 0.54 [0.47, 0.59] | -0.06 [-0.12, -0.01] | 0.64 [0.61, 0.71] | 0.0 [-0.06, 0.04] |
| **Fibroblasts (N = 615 cells)** | 0.34 [0.26, 0.4] | -0.04 [-0.1, 0.01] | 0.35 [0.3, 0.39] | -0.04 [-0.09, 0.0] | 0.36 [0.33, 0.42] | -0.04 [-0.09, -0.01] |
| **Lymphocytes (N = 1,962 cells)** | 0.72 [0.65, 0.74] | 0.17 [0.12, 0.21] | 0.47 [0.42, 0.51] | -0.09 [-0.12, -0.05] | 0.56 [0.53, 0.61] | 0.04 [0.01, 0.06] |
| **Macrophages (N = 414 cells)** | 0.25 [0.17, 0.31] | 0.14 [0.06, 0.2] | 0.13 [0.09, 0.15] | 0.0 [-0.03, 0.04] | 0.28 [0.26, 0.33] | 0.06 [-0.01, 0.1] |
| **Plasma Cells (N = 438 cells)** | 0.62 [0.51, 0.67] | 0.17 [0.11, 0.23] | 0.3 [0.23, 0.35] | -0.18 [-0.23, -0.11] | 0.47 [0.44, 0.54] | -0.06 [-0.13, -0.02] |

| | STAD | | | | | |
|---|---|---|---|---|---|---|
| | **Recall** | | **Precision** | | **F1 Score** | |
| **Cell Class** | **Model vs. Avg Pathologist [95% CI]** | **Δrecall [95% CI]** | **Model vs. Avg Pathologist [95% CI]** | **ΔPrecision [95% CI]** | **Model vs. Avg Pathologist [95% CI]** | **ΔF1 [95% CI]** |
| **Cancer Cells (N = 2,480 cells)** | 0.67 [0.63, 0.72] | -0.06 [-0.12, 0.0] | 0.62 [0.56, 0.67] | -0.11 [-0.16, -0.07] | 0.64 [0.59, 0.67] | -0.08 [-0.12, -0.05] |
| **Fibroblasts (N = 1,667 cells)** | 0.27 [0.23, 0.3] | -0.17 [-0.21, -0.13] | 0.59 [0.54, 0.64] | 0.15 [0.1, 0.2] | 0.35 [0.3, 0.39] | -0.04 [-0.08, -0.01] |
| **Lymphocytes (N = 3,493 cells)** | 0.68 [0.63, 0.71] | -0.02 [-0.05, 0.0] | 0.66 [0.62, 0.7] | -0.04 [-0.07, -0.01] | 0.66 [0.61, 0.69] | -0.02 [-0.03, 0.0] |
| **Macrophages (N = 1,209 cells)** | 0.08 [0.06, 0.09] | -0.06 [-0.13, 0.02] | 0.25 [0.2, 0.31] | 0.12 [0.04, 0.22] | 0.11 [0.1, 0.15] | 0.03 [-0.04, 0.06] |
| **Plasma Cells (N = 636 cells)** | 0.54 [0.47, 0.58] | -0.06 [-0.12, -0.02] | 0.54 [0.44, 0.6] | -0.07 [-0.14, -0.01] | 0.5 [0.42, 0.55] | -0.02 [-0.07, 0.02] |
