## Supplementary Table S5 for "Spatial mapping of immunosuppressive CAF gene signatures in H&E-stained images using additive multiple instance learning"

**Supplementary Table 5. Summary table of cell model performance characteristics.**

**BLCA**

| Tissue Region | Precision | | Recall | | F1 Score | |
|---|---|---|---|---|---|---|
| | Model vs. Avg Pathologist [95% CI] | Mean difference [95% CI] | Model vs. Avg Pathologist [95% CI] | Mean difference [95% CI] | Model vs. Avg Pathologist [95% CI] | Mean difference [95% CI] |
| **Cancer (N = 111,425 pixels)** | 0.63 [0.57, 0.69] | -0.04 [-0.08, 0.0] | 0.71 [0.64, 0.75] | 0.04 [-0.02, 0.09] | 0.68 [0.62, 0.72] | 0.0 [-0.03, 0.05] |
| **Cancer Stroma (N = 78,895 pixels)** | 0.5 [0.42, 0.54] | -0.03 [-0.09, 0.0] | 0.52 [0.44, 0.6] | 0.01 [-0.05, 0.09] | 0.5 [0.43, 0.57] | 0.0 [-0.06, 0.04] |
| **Necrosis (N = 51,484 pixels)** | 0.38 [0.42, 0.24] | -0.19 [-0.32, -0.12] | 0.64 [0.5, 0.71] | 0.13 [0.04, 0.25] | 0.61 [0.48, 0.68] | -0.04 [-0.17, 0.05] |
| **Normal Tissue (N = 187,110 pixels)** | 0.63 [0.55, 0.67] | -0.04 [-0.11, -0.01]] | 0.73 [0.65, 0.77] | 0.04 [-0.02, 0.1] | 0.67 [0.61, 0.72] | 0.02 [-0.06, 0.05] |

**STAD**

| Tissue Region | Precision | | Recall | | F1 Score | |
|---|---|---|---|---|---|---|
| | Model vs. Avg Pathologist [95% CI] | Mean difference [95% CI] | Model vs. Avg Pathologist [95% CI] | Mean difference [95% CI] | Model vs. Avg Pathologist [95% CI] | Mean difference [95% CI] |
| **Cancer (N = 112,373 pixels)** | 0.73 [0.67, 0.77] | -0.04 [-0.08, -0.02] | 0.77 [0.71, 0.82] | -0.0 [-0.05, 0.04] | 0.74 [0.69, 0.78] | -0.02 [-0.04, 0.01] |
| **Cancer Stroma (N = 103,702 pixels)** | 0.58 [0.5, 0.66] | -0.03 [-0.09, 0.01] | 0.62 [0.55, 0.67] | -0.0 [-0.06, 0.06] | 0.59 [0.51, 0.65] | -0.01 [-0.05, 0.03] |
| **Necrosis (N = 31,256 pixels)** | 0.47 [0.3, 0.61] | -0.17 [-0.25, -0.07] | 0.66 [0.48, 0.78] | 0.03 [-0.09, 0.18] | 0.55 [0.39, 0.67] | -0.08 [-0.17, 0.02] |
| **Mucin (N = 19,468 pixels)** | 0.42 [0.24, 0.54] | -0.14 [-0.27, -0.06] | 0.63 [0.47, 0.75] | 0.08 [-0.01, 0.19] | 0.51 [0.31, 0.6] | -0.01 [-0.13, 0.05] |
| **Normal Tissue (N = 92,359 pixels)** | 0.63 [0.53, 0.7] | -0.09 [-0.16, -0.04] | 0.66 [0.58, 0.73] | -0.06 [-0.13, 0.03] | 0.64 [0.56, 0.69] | -0.07 [-012, -0.02]. |

**PRAD**

| Tissue Region | Precision | | Recall | | F1 Score | |
|---|---|---|---|---|---|---|
| | Model vs. Avg Pathologist [95% CI] | Mean difference [95% CI] | Model vs. Avg Pathologist [95% CI] | Mean difference [95% CI] | Model vs. Avg Pathologist [95% CI] | Mean difference [95% CI] |
| **Cancer (N = 99,378 pixels)** | 0.776 [0.711, 0.812] | -0.001 [-0.042, 0.019] | 0.748 [0.700, 0.792] | -0.056 [-0.107, -0.011] | 0.757 [0.699, 0.793] | -0.011 [0.013, -0.039] |
| **Cancer Stroma (N = 90,984 pixels)** | 0.578 [0.504, 0.636] | -0.119 [-0.182, -0.078] | 0.757 [0.724, 0.784] | -0.108 [-0.235, 0.184] | 0.649 [0.590, 0.692] | -0.034 [0.001, -0.080] |
| **Necrosis (N = 12,839 pixels)** | 0.457 [0.316, 0.542] | -0.167 [-0.279, -0.085] | 0.501 [0.345, 0.743] | 0.061 [0.021, 0.100] | 0.447 [0.321, 0.599] | -0.125 [0.009, -0.288] |
| **Normal Tissue (N = 141,394 pixels)** | 0.741 [0.661, 0.797] | -0.051 [-0.109, -0.012] | 0.735 [0.670, 0.791] | -0.029 [-0.064, 0.005] | 0.730 [0.662, 0.786] | -0.049 [-0.017, -0.089] |
