## Supplementary Figures 1-11 for "Spatial mapping of immunosuppressive CAF gene signatures in H&E-stained images using additive multiple instance learning"

### Supplemental Figures

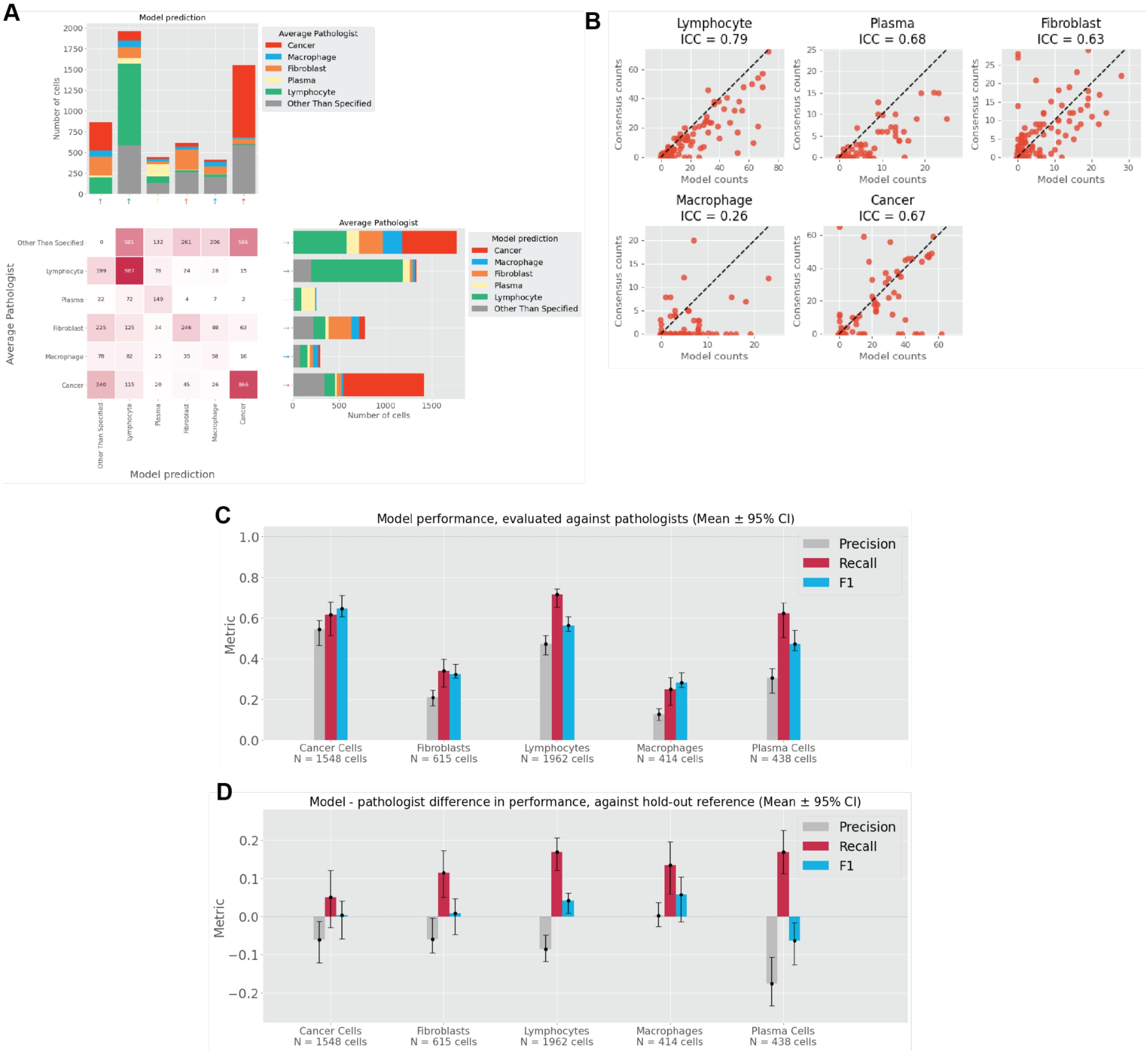


**Supplementary Figure S1. Performance of cell classification model in bladder cancer.** A) Comparison of model-predicted cell types to average pathologist annotations. The bar graph at the top left depicts the breakdown of average pathologist annotations for each class of model prediction (precision). The bar graph at the bottom right shows the breakdown of model predictions for each class of average pathologist annotation (recall). “Other Than Specified” refers to predictions of classes other than those listed, or background class. B) Agreement between model-derived cell counts and pathologist consensus counts . C) Precision, Recall, and F1 scores of model predictions compared to pathologists’ annotations in nested pairwise fashion[19]. D) Difference in the nested pairwise metric comparing mean difference between model and individual pathologist performance. Positive values indicate that the model out-performed pathologists when evaluated against held-out pathologists, while negative values indicate the model under-performed pathologists. Confidence intervals were obtained by bootstrapping.

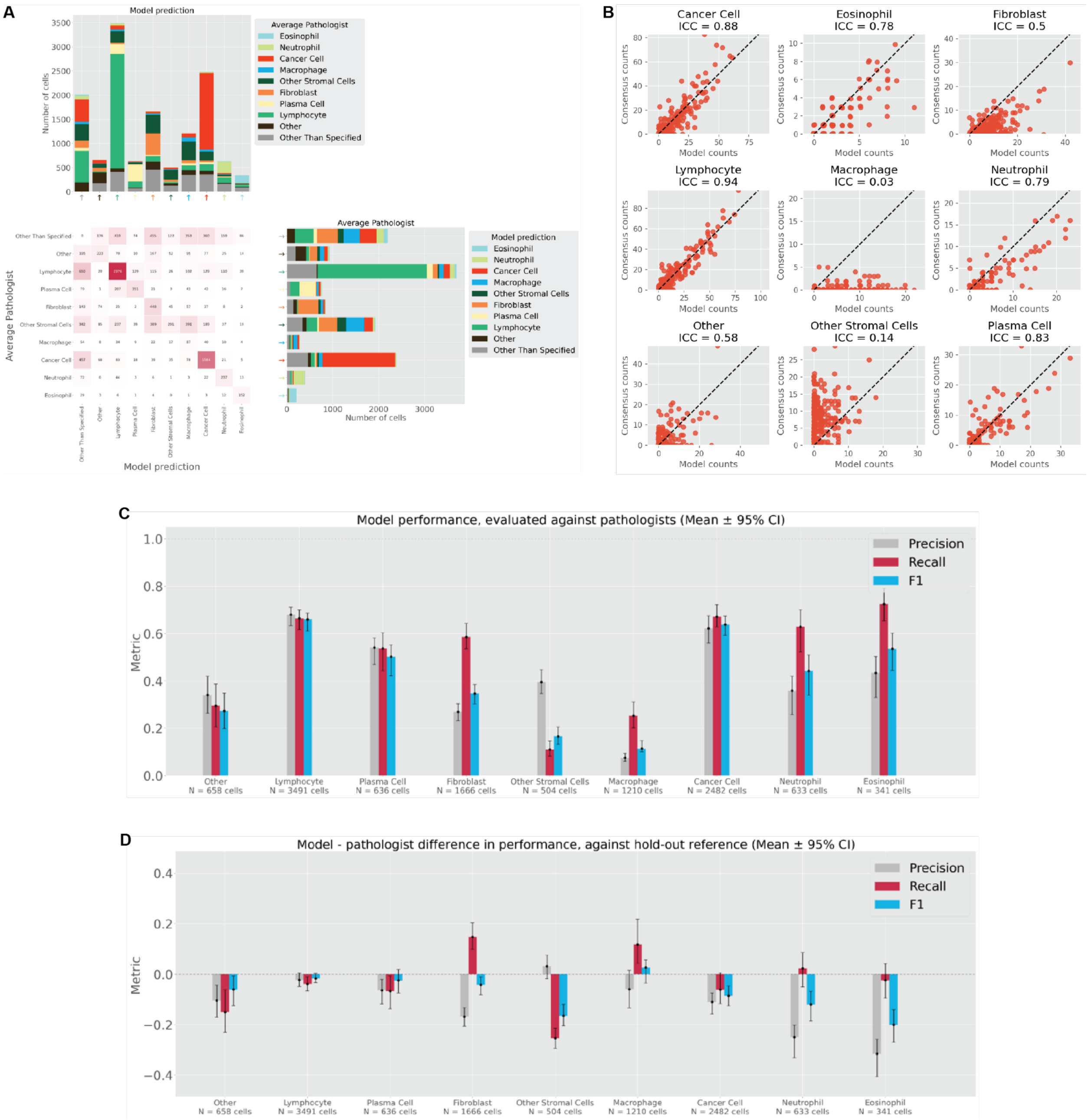


**Supplementary Figure S2. Performance of cell classification model in gastric cancer.** A) Comparison of model-predicted cell types to average pathologist annotations. The bar graph at the top left depicts the breakdown of average pathologist annotations for each class of model prediction (precision). The bar graph at the bottom right shows the breakdown of model predictions for each class of average pathologist annotation (recall). “Other Than Specified” refers to predictions of classes other than those listed, or background class. B) Agreement between model-derived cell counts and pathologist consensus counts . C) Precision, Recall, and F1 scores of model predictions compared to pathologists’ annotations in nested pairwise fashion[19]. D) Difference in the nested pairwise metric comparing mean difference between model and individual pathologist performance. Positive values indicate that the model out-performed pathologists when evaluated against held-out pathologists, while negative values indicate the model under-performed pathologists. Confidence intervals were obtained by bootstrapping.

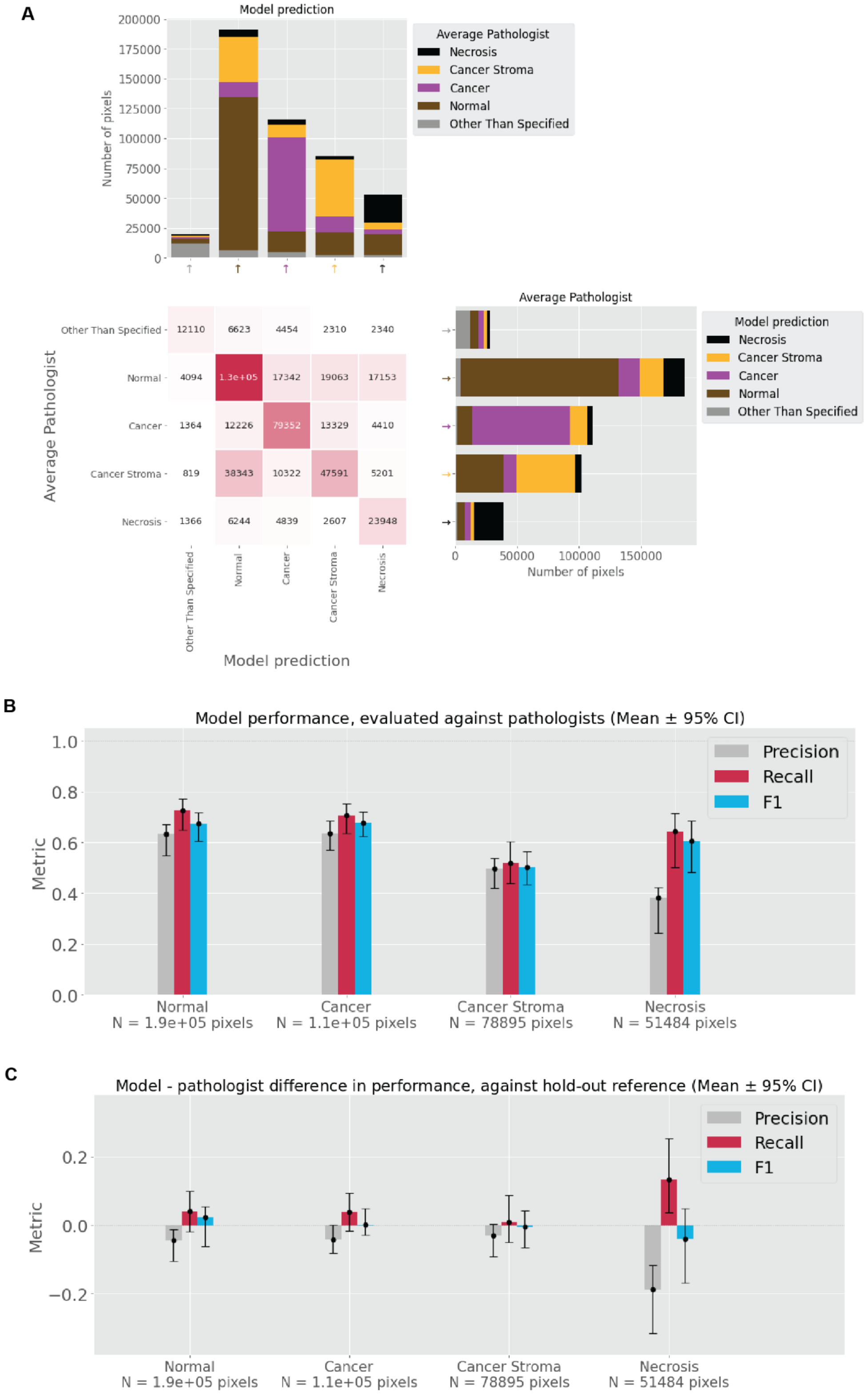


**Supplementary Figure S3. Performance of tissue classification model in bladder cancer.** A) Confusion matrix of tissue model classification compared to average pathologist prediction. The bar graph at the top left depicts the breakdown of average pathologist annotations for each class of model prediction (precision). The bar graph at the bottom right shows the breakdown of model predictions for each class of average pathologist annotation (recall). “Other Than Specified” refers to predictions of classes other than those listed, or background class. B) Precision, Recall, and F1 scores of model predictions compared to pathologists’ annotations in nested pairwise fashion[19]. C) Difference in the nested pairwise metric comparing mean difference between model and individual pathologist performance. Positive values indicate that the model out-performed pathologists when evaluated against held-out pathologists, while negative values indicate the model under-performed pathologists. Confidence intervals were obtained by bootstrapping.

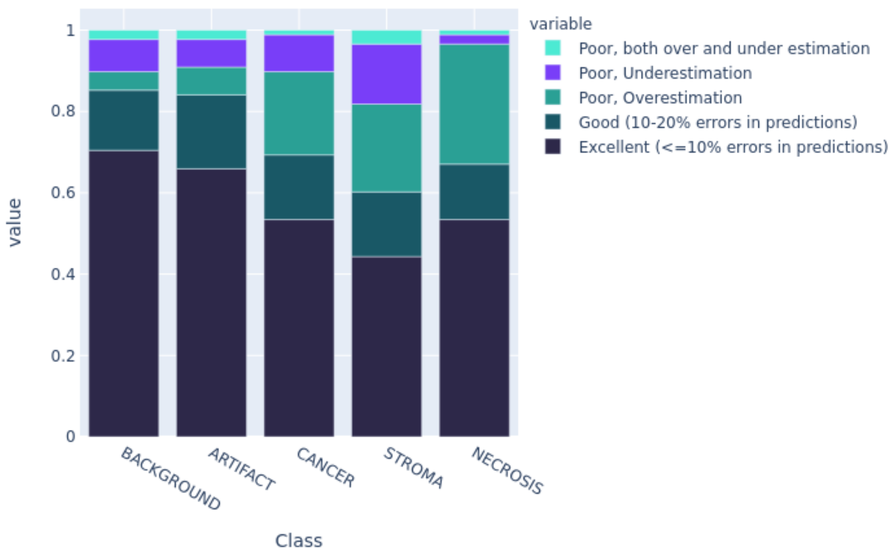


**Supplementary Figure S4. Performance of tissue classification model in breast cancer.** The percent (%) of WSIs out of 88 WSIs in the evaluation dataset per pathologist evaluator assessment of model performance. Colors correspond to the pathologist evaluator's assessment (<10% error, 10-20% error, or >20% error) and failure mode (overestimation, underestimation, or both).

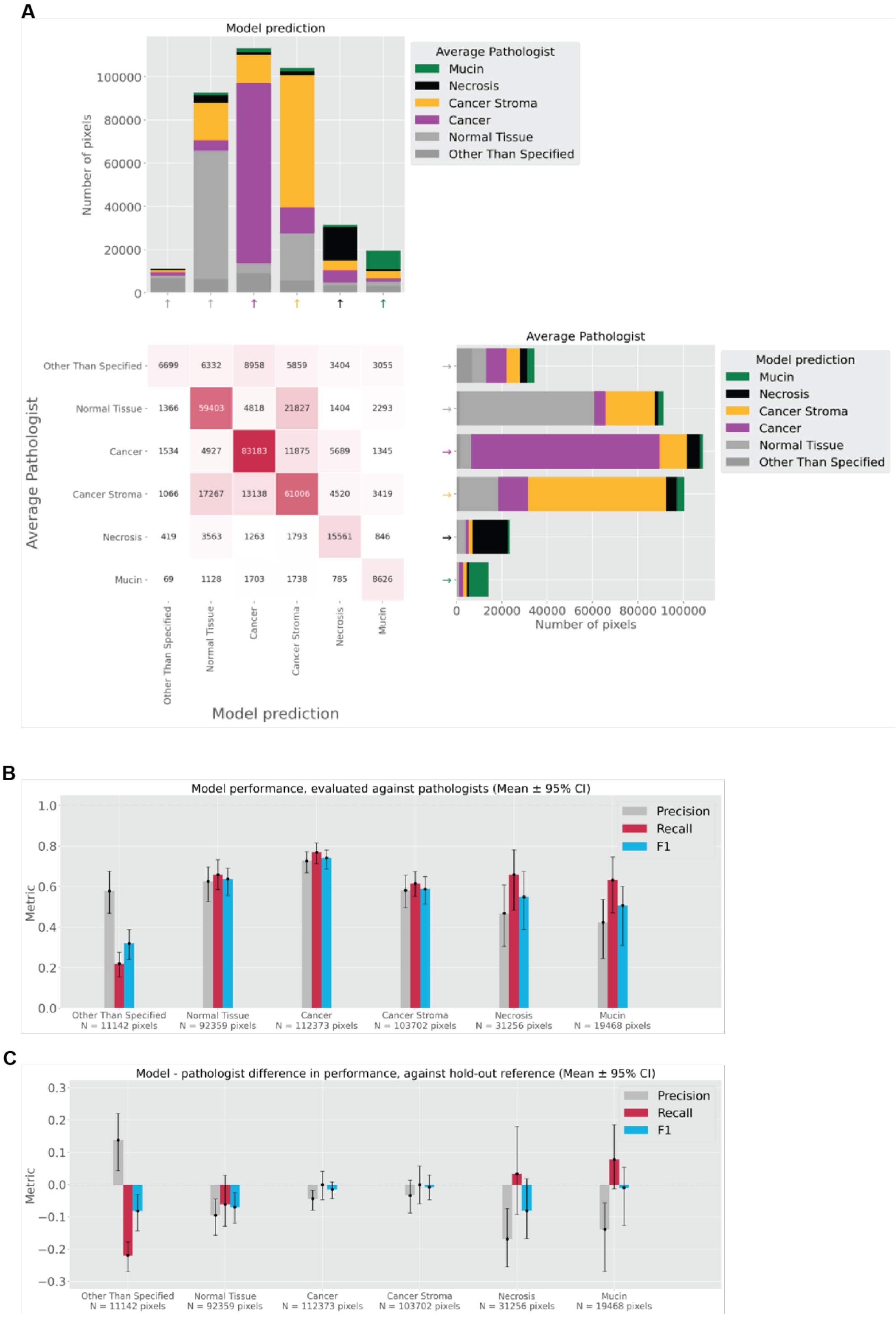


**Supplementary Figure S5. Performance of tissue classification model in gastric cancer.** A) Confusion matrix of tissue model classification compared to average pathologist prediction. The bar graph at the top left depicts the breakdown of average pathologist annotations for each class of model prediction (precision). The bar graph at the bottom right shows the breakdown of model predictions for each class of average pathologist annotation (recall). “Other Than Specified” refers to predictions of classes other than those listed, or background class. B) Precision, Recall, and F1 scores of model predictions compared to pathologists’ annotations in nested pairwise fashion[19]. C) Difference in the nested pairwise metric comparing mean difference between model and individual pathologist performance. Positive values indicate that the model out-performed pathologists when evaluated against held-out pathologists, while negative values indicate the model under-performed pathologists. Confidence intervals were obtained by bootstrapping.

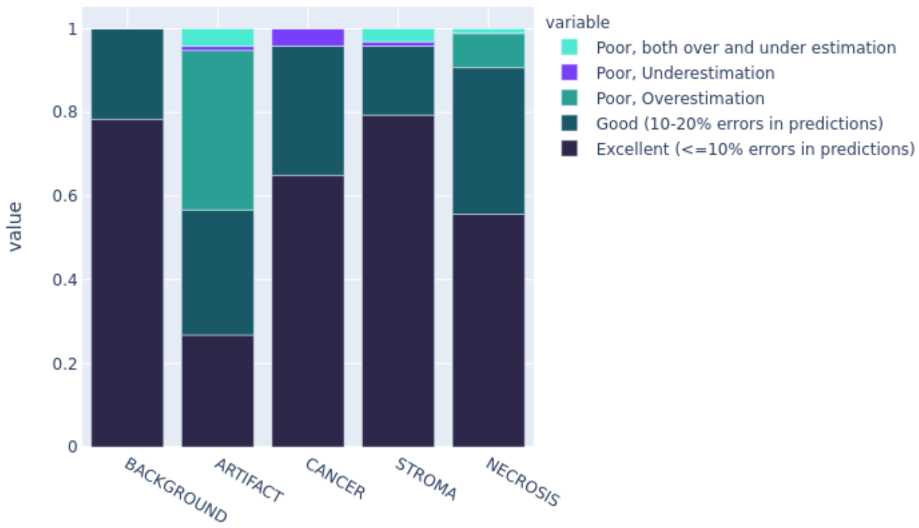


**Supplementary Figure S6. Performance of tissue classification model in NSCLC.** Percent of WSI (out of 97 test WSIs) that passed a qualitative pathologist evaluation. The main mode of failure is highlighted for cases with poor performance.

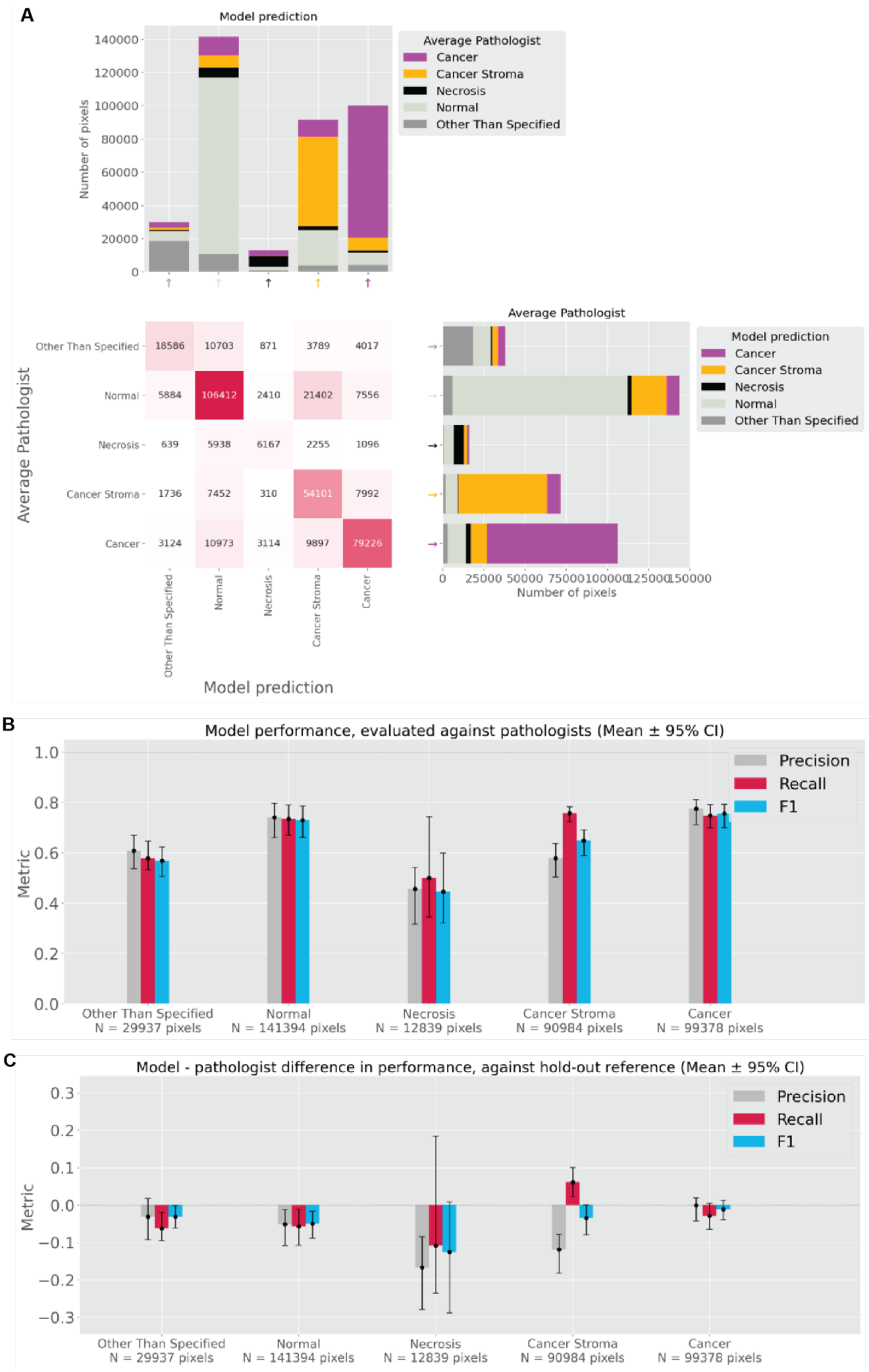


**Supplementary Figure S7. Performance of tissue classification model in prostate cancer.** A) Confusion matrix of tissue model classification compared to average pathologist prediction. The bar graph at the top left depicts the breakdown of average pathologist annotations for each class of model prediction (precision). The bar graph at the bottom right shows the breakdown of model predictions for each class of average pathologist annotation (recall). “Other Than Specified” refers to predictions of classes other than those listed, or background class. B) Precision, Recall, and F1 scores of model predictions compared to pathologists’ annotations in nested pairwise fashion[19]. C) Difference in the nested pairwise metric comparing mean difference between model and individual pathologist performance. Positive values indicate that the model out-performed pathologists when evaluated against held-out pathologists, while negative values indicate the model under-performed pathologists. Confidence intervals were obtained by bootstrapping.

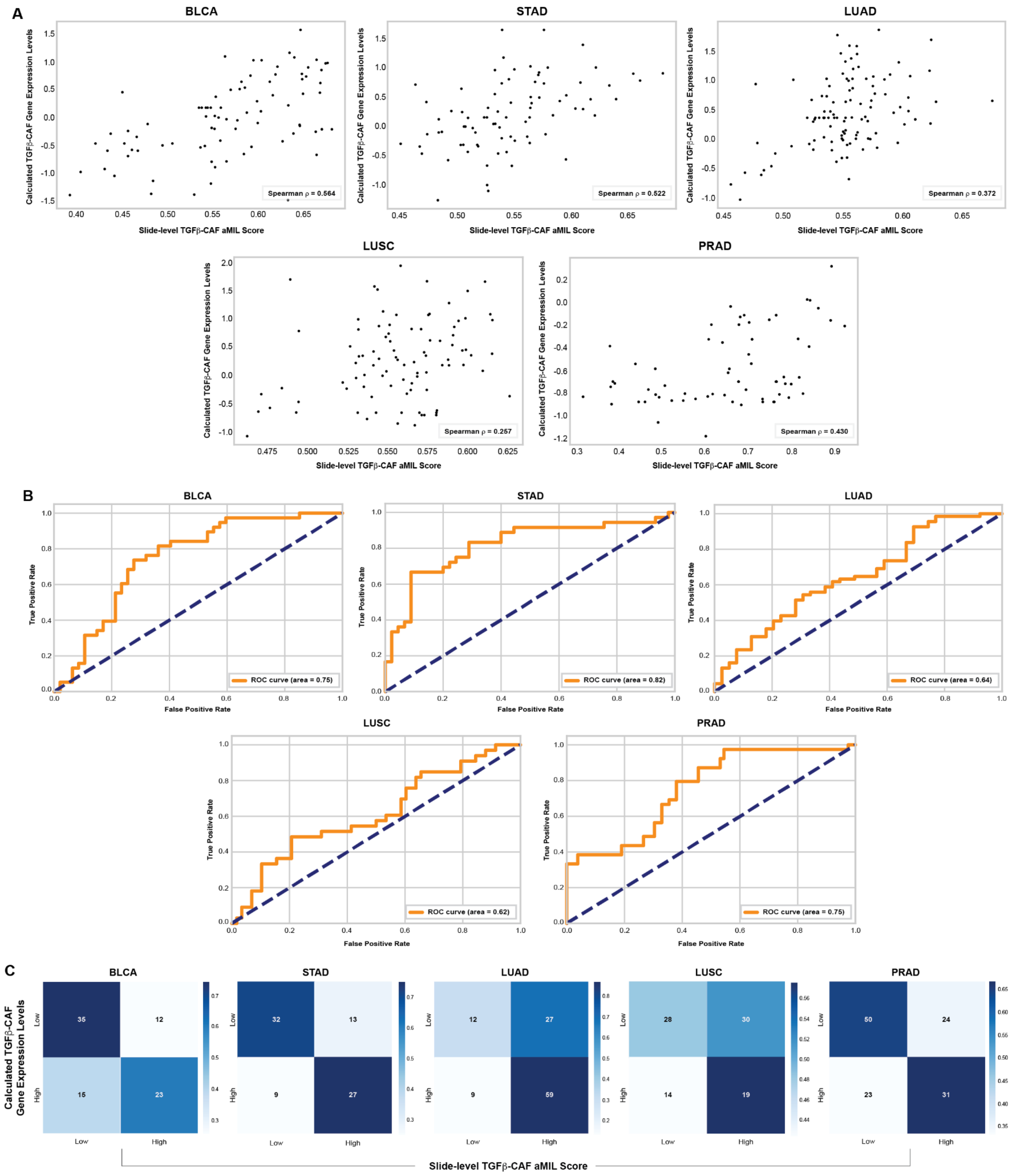


**Supplementary Figure S8. Performance of models in predicting TGFβ-CAF levels in bladder, lung, stomach, and prostate cancer.** A) Correlation of model-predicted slide-level TGFβ-CAF scores with levels calculated from sequencing data for each case. B) Performance of aMIL model in predicting TGFβ-CAF levels. C) Confusion matrix depicting agreement of model predictions with ground truth. The number of slides in each category is indicated. All analyses were performed in the test sets for each indication.

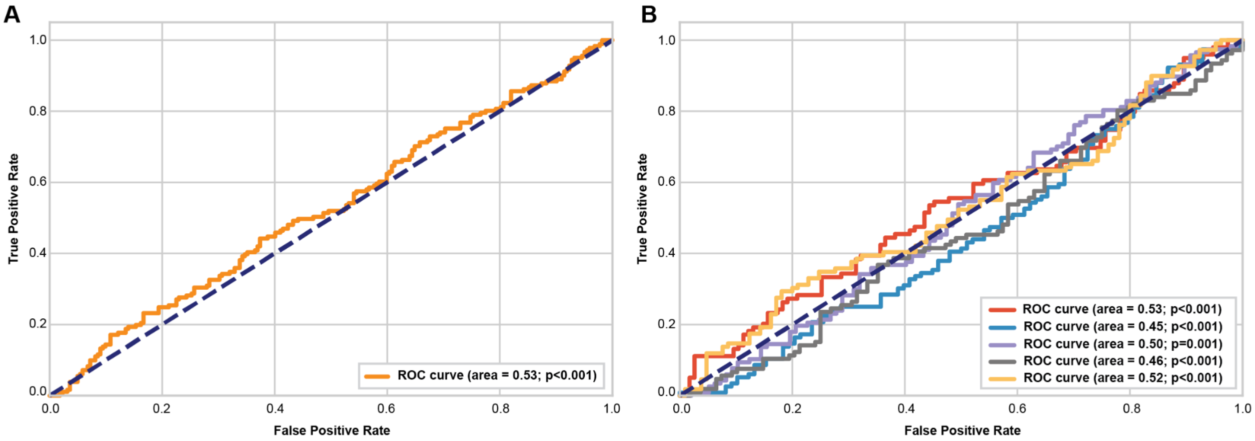


**Supplementary Figure S9. Performance of negative control models in predicting TGFβ-CAF levels.** A) ROC curve for model trained on slide background. B) ROC curve for five models trained using random labels.

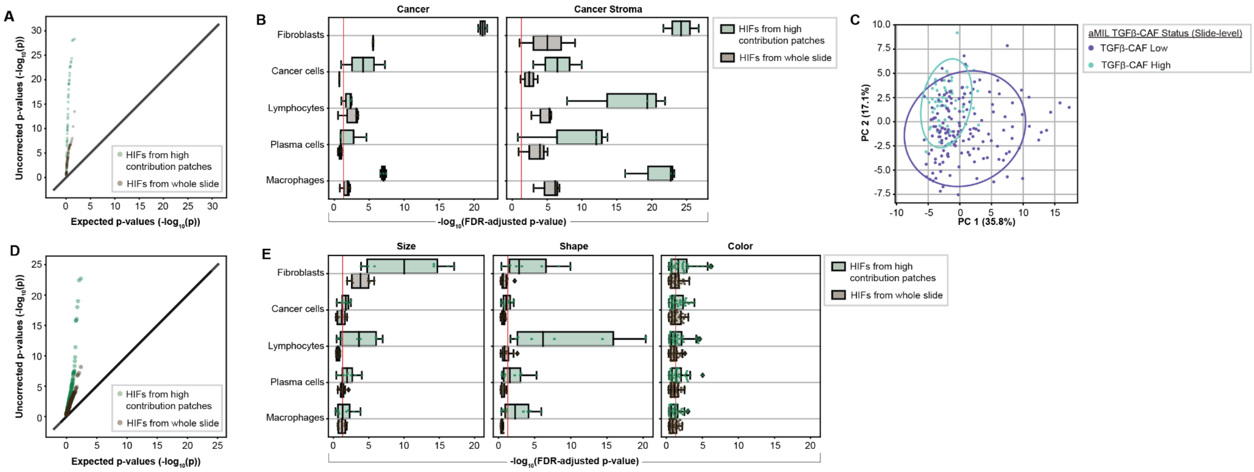


**Supplementary Figure S10. Features extracted from high-contribution patches vs. entire slide.** A) QQ plot of p values of cell and tissue features derived from high contribution regions (green) or from whole slides (gray). Each point represents a unique feature. B) Quantification of cell features derived from high contribution patches (green) or whole slides (gray). C) Principal components analysis using HIFs extracted from whole slides. A projection of patches onto the first two principal components, colored by predicted TGFβ-CAF status, is shown. D) QQ plot of p values of nuclear features derived from high contribution regions (green) or from whole slides (gray). Each point represents a unique feature. E) Quantification of nuclear features derived from high contribution patches (green) or whole slides (gray).

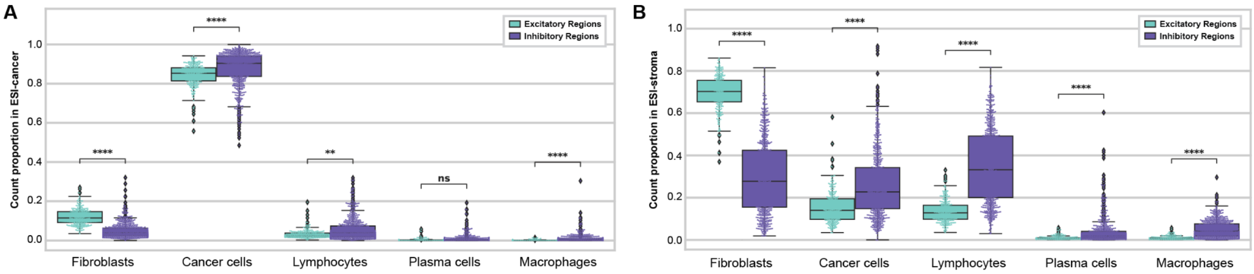


**Supplemental Figure S11. Quantification of relative cell numbers within 120 μm of epithelial-stromal interface.** Densities of predicted cell types within 120 μm of the cancer-stroma boundary are shown A) within the cancer and B) within the stroma.
